# TLR4 regulates proinflammatory intestinal immune responses mediated by an atopic gut microbiota

**DOI:** 10.1101/2022.07.21.501027

**Authors:** Evelyn Campbell, Lisa Maccio-Maretto, Lauren A. Hesser, Andrea M. Kemter, Roberto Berni Canani, Rita Nocerino, Lorella Paparo, Robert T. Patry, Cathryn R. Nagler

## Abstract

The increasing prevalence of food allergies has been causally associated with the depletion of allergy protective intestinal bacteria. However, few studies have investigated the role of the gut microbiota in promoting allergic responses. In a cohort of infants affected by cow’s milk allergy (CMA), we have identified a patient with a proinflammatory and atopic microbiota. In comparison to a healthy microbiota, this CMA-associated gut microbiota has increased abundance of Bacteroidetes, a Gram-negative phylum of bacteria that has been associated with increased incidence of allergy. Using this microbiota, we investigated the host-microbe interactions that mediate these intestinal inflammatory responses. To examine these interactions, we used mice with global and conditional abrogation in TLR4 signaling, since Gram-negative bacteria signal through this receptor via membrane-derived lipopolysaccharide (LPS). We show that this donor’s microbiota induces expression of serum amyloid A1 (*Saa1*) and other Th17-, B cell-, and Th2-associated genes in the ileal epithelium. Accordingly, this microbiota also induces Th17 cells, as well as regulatory T cell populations and fecal IgA. Importantly, we used both antibiotic treated SPF and rederived germ-free mice with a conditional mutation of TLR4 in the CD11c^+^ compartment to demonstrate that the induction of proinflammatory genes, fecal IgA, and Th17 cells is dependent on TLR4 signaling. Furthermore, metagenomic sequencing revealed that the CMA-associated gut microbiota also has increased abundance of LPS biosynthesis genes. Lastly, upon sensitization with β-lactoglobulin, this CMA microbiota induces a TLR4-dependent mixed type 2/type 3 response in innate lymphoid cells (ILCs) during the early phases of allergic sensitization. Taken together, our results show that a Bacteroidetes-enriched microbiota with increased abundance of LPS genes promotes proinflammatory gene expression and a mixed type 2/type 3 response in a subset of infants with cow’s milk allergy.

**Paper Highlights:**

1. A cow’s milk allergy (CMA)-associated gut microbiota has an enrichment of Bacteroidetes, which is associated with atopy
2. The CMA-associated gut microbiota promotes intestinal inflammation, which includes inflammatory gene expression, induction of Th17 cells, and production of IgA
3. Proinflammatory responses induced by the CMA-associated gut microbiota are dependent on TLR4 signaling in various cellular compartments
4. Upon sensitization, the CMA-associated gut microbiota induces an innate mixed type 2/type 3 inflammatory response

## Introduction

Within the last century, food allergies (FAs), along with asthma, eczema, and other atopic diseases, have become more common within the United States and other first-world countries (Gupta et al., 2018; Brough et al., 2021). FAs can present with a multitude of symptoms, such as hives, itchy throat, wheezing, difficulty breathing, vomiting, and life-threatening anaphylactic shock. As therapeutic interventions remain limited, FAs are becoming a growing global health concern (Iweala and Nagler, 2019; Warren et al., 2022). Concurrent with this generational increase in FAs, Western lifestyle factors, such as poor dietary habits, misuse of antibiotics, and increased Cesarean birth, have contributed to a disruption in gut homeostasis and dysbiosis – a collective disturbance of the microbiota (Renz and Skevaki, 2021). Recent findings from our group and others have demonstrated that the microbiota of allergic children is more dysbiotic than their healthy counterparts (De Filippis et al., 2021; Berni Canani et al., 2018; Bao et al., 2021). Numerous epidemiological studies report direct correlations between juvenile microbial perturbation and atopic diseases (Marra et al., 2009; Russell et al., 2012; Metsälä et al., 2012), adding to increasing evidence that early life immunological education is regulated by microbial exposure and can impact susceptibility to inflammatory diseases later in life (Knoop et al., 2017; Al Nabhini et al., 2019). Therefore, it is critical that we understand the host-microbial interactions contributing to the development of allergic immune responses to innocuous food antigens.

Recent studies have started to elucidate the mechanisms by which protective members of the microbiota regulate food allergic responses. We and others have shown that Clostridia species, such as *Lachnospiraceae,* prevent allergic responses to food antigens in murine models (Stefka et al., 2014; Feehley et al., 2019; Abdel-Gadir et al., 2019). Bacterial products, such as short chain fatty acids (Tan et al., 2016; Paparo and Nocerino et al., 2020; Wang and Cao et al., 2022), Toll-like receptor (TLR) ligands (Shim et al., 2016), and AhR ligands (Bae et al., 2016) can protect from atopic diseases and are produced by Clostridial species. Studies have also shown that microbially-derived products can exacerbate type 2 inflammation (Zhang et al., 2017; Lee et al., 2017); however, the underlying mechanisms behind these allergic responses, particularly to food antigens, remain poorly understood.

Innate immune signaling through TLRs is important for initiating adaptive humoral and cellular responses in allergic diseases, such as food allergies and asthma (Bashir et al., 2004; Matsushita and Yoshimoto, 2014; Penders et al., 2010). Host signaling through Toll-like receptor 4 (TLR4) has long been known to regulate type 2 immune responses in models of these diseases (Bashir et al., 2004; Eisenbarth et al., 2002). The context in which lipopolysaccharide (LPS), a bacterial agonist of TLR4, is recognized by the host can dictate whether protective or proallergic immune responses are induced. Low dose exposure of LPS drives type 2 inflammation in mouse models of allergic asthma, while high doses are protective (Eisenbarth et al., 2002; Radermecker et al., 2019; Bachus et al., 2019). Thus, the context of TLR4 stimulation and source of LPS can regulate sensitivity to food allergens.

Here, we investigate the role of TLR4 in regulating allergic sensitization in the context of the microbiota from a severely cow’s milk allergic (CMA) infant. We show that this CMA microbiota induces expression of epithelial serum amyloid A1 (*Saa1*). We demonstrate that the induction of *Saa1,* Tregs, and Th17 cell populations in the ileum are regulated by TLR4 signaling, particularly in CD11c^+^ cells. Analyzing the microbiome of mice colonized with this CMA infant microbiota revealed an increased representation of Gram-negative bacteria, particularly Bacteroidetes, when compared to mice colonized with the microbiota of a healthy infant. Finally, we show that after sensitization with a cow’s milk allergen, the CMA microbiota induces a mixed type 2/type 3 response in ILCs that is regulated by TLR4. These results illuminate a previously unappreciated role for the TLR4-LPS innate signaling axis as a mechanism by which the CMA-associated dysbiotic gut microbiota could contribute to inflammatory responses associated with FA.

## Results

### The microbiota from a severely symptomatic CMA donor promotes an intestinal inflammatory response

Previous work from our lab examined the role of the microbiota from a cohort of healthy and IgE-mediated cow’s milk allergy (CMA) infants to understand microbial contributors to FA (Feehley et al., 2019). Members of the healthy infant gut microbiota, particularly *Anaerostipes caccae*, were able to protect against allergic sensitization to the major cow’s milk allergen β- lactoglobulin (BLG). The CMA-associated gut microbiota was unable to protect against allergic sensitization to BLG and elicited an elevated BLG-specific response compared to germ-free (GF) and healthy-colonized mice, suggesting that the CMA-associated gut microbiota may promote allergic responses (Feehley et al., 2019). In the present study, we focused on one CMA infant donor that exhibited more severe symptoms upon consumption of cow’s milk compared to other CMA infant donors within this cohort. This donor, CMA Donor 5, experienced urticaria and gastrointestinal symptoms within a few minutes after feeding with cow’s milk. Moreover, this donor presented with gastrointestinal inflammation, as observed by nodular hyperplasia on the intestinal walls of the rectum. Histological evaluation revealed eosinophilic infiltration within the gastrointestinal tissue, supporting an immunological basis for the symptoms experienced by the patient (**Figure 1A**).

**Figure 1.**
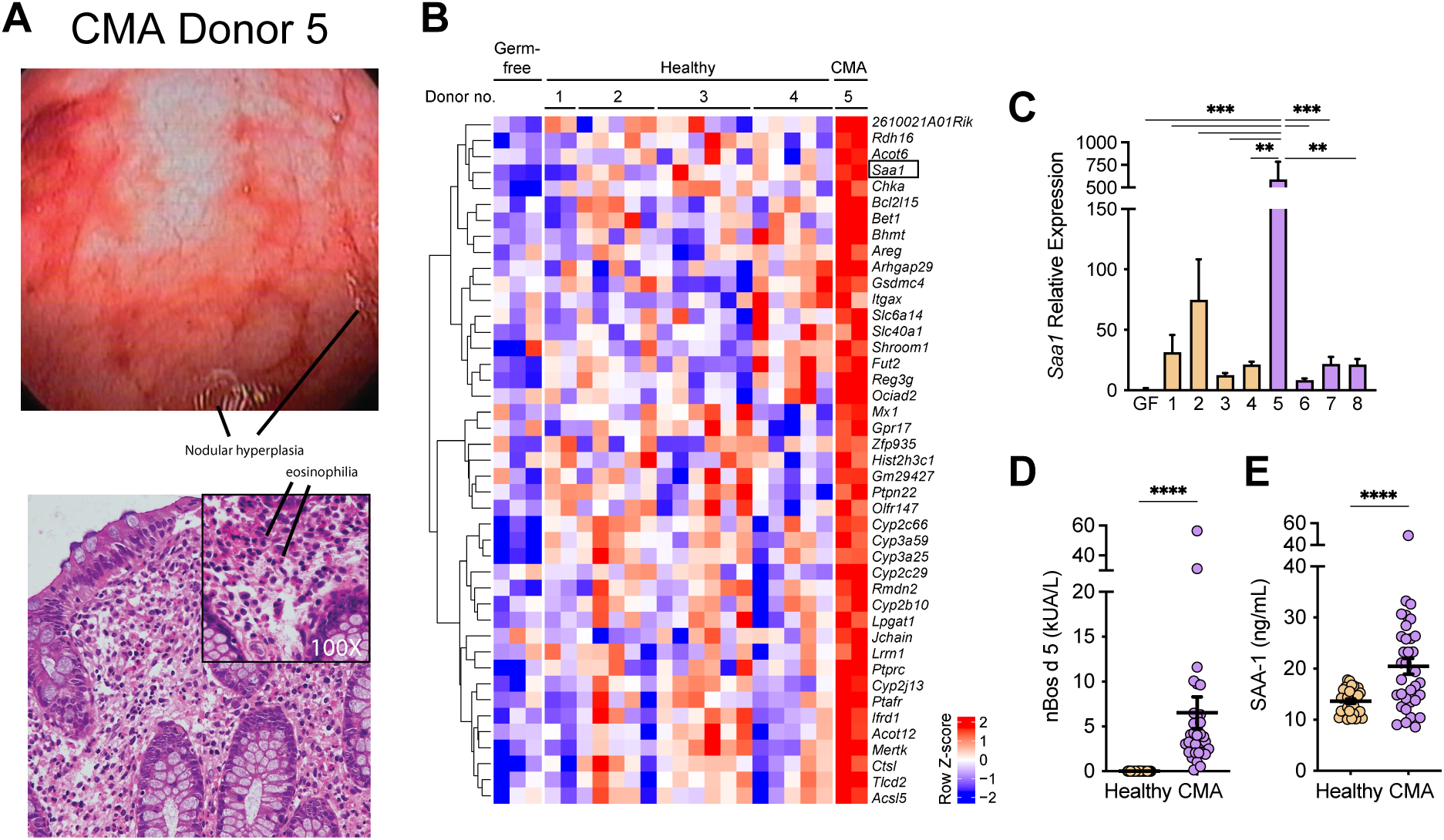
TLR4 regulates expression of Th17-associated genes in the ileal epithelium of Donor 5-colonized mic. A) Colonoscopy and histology images of the GI tract of CMA Donor 5 after consumption of cow’s milk. B) Heatmap of relative abundance of gene transcripts significantly upregulated in CMA Donor 5 colonized mice compared to GF and mice colonized with healthy donors 1 – 4. C) qRT-PCR analysis of *Saa1* relative expression in GF mice and mice colonized with the microbiota from healthy donors (donors 1-4) and CMA donors (5-8). D) Serum concentrations of IgE specific for the nBos d 5 epitope of BLG and E) SAA-1 from age and sex-matched pediatric healthy and CMA patients. Symbols represent individual patients, bars represent mean ±S.E.M, n=30 patients per group. For B, significance was determined through Benjamini-Hochberg multiple testing correction, with a FDR ≤ 0.1. For C, 2-way ANOVA with multiple testing using the Benjamini, Krieger, and Yekutili method was performed. For D-E, Wilcoxon rank sum test was performed between both conditions. \*\**P* < 0.01, \*\*\**P* < 0.001, \*\*\*\**P* < 0.0001

Stool samples from CMA Donor 5 were used to colonize GF mice to examine the role of the microbiota in mediating allergic sensitization to BLG. In response to the initial colonization with the CMA Donor 5 microbiota, some mice appeared to be lethargic and exhibited diarrhea and crusted fecal matter around the anus. After euthanasia of mice colonized with the CMA Donor 5 microbiota for one week, some mice were observed to have bloody intestinal contents in the ileum and loose stool pellets in the colon (**Figure S1**). In the original publication (Feehley et al., 2019), ileal and colonic tissue was collected from mice colonized with feces from each of the eight infant donors at sacrifice after sensitization with BLG plus cholera toxin (CT) and at 5-6 months post colonization. The tissues were fixed in formalin and stained with hematoxylin and eosin (H & E) or placed in Carnoy’s fixative for periodic acid-Schiff (PAS) staining. A gastrointestinal pathologist examined the stained sections in a blinded fashion and found no evidence of pathology, ruling out an infectious etiology (Feehley et al., 2019, Extended Data Figs. 1 and 2). The similarity of the symptomatic manifestations observed in both CMA Donor 5 and in mice colonized with the Donor 5 microbiota led to the hypothesis that this microbiota may be contributing to allergic responses by inducing transient intestinal inflammation.

**Figure 2:**
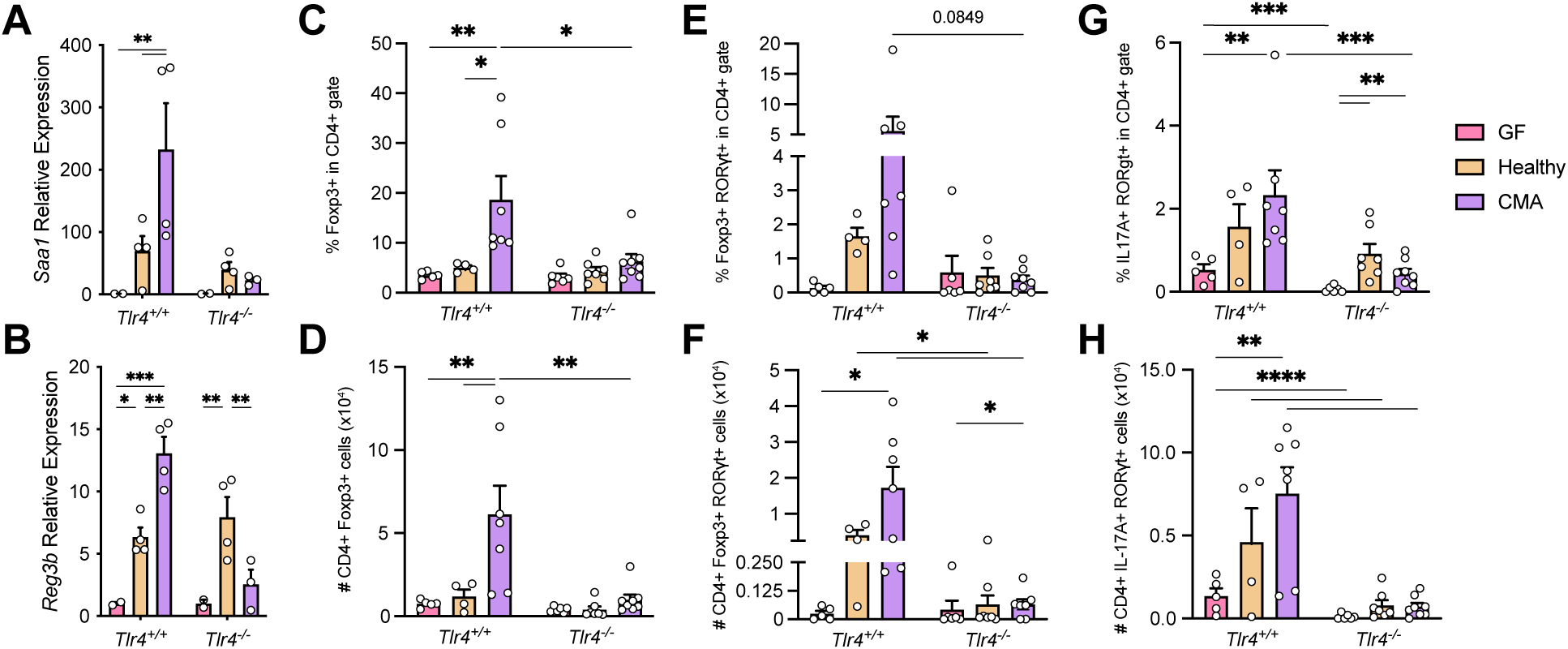
TLR4 regulates T cell populations in the ileum of Donor 5-colonized mice. A) Relative expression of *Saa1* and B) *Reg3b* in ileal ECs from GF, healthy- (Donor 1) or CMA- colonized (Donor 5) *Tlr4^+/+^* and *Tlr4^-/-^* mice one-week post-colonization. C-H, Frequency, and absolute numbers of C-D) Foxp3^+^ (SP) Tregs, E-F) Foxp3^+^ RORγt^+^ (DP) Tregs, and G-H) IL-17^+^ RORγt^+^ Th17 cells in the ileal lamina propria of *Tlr4^+/+^* and *Tlr4^-/-^* GF, healthy-colonized, and CMA-colonized mice. Data are pooled from at least two independent experiments per condition. Statistics were calculated using 2-way ANOVA with multiple testing using the Benjamini, Krieger, and Yekutili method. \**P* < 0.05, \*\**P* < 0.01, \*\*\**P* < 0.001, \*\*\*\**P* < 0.0001

In the context of FAs and other atopic diseases, the epithelium acts not only as a barrier between exogenous antigens and the immune system, but also receives signals from these exogenous antigens through innate receptors and relays information about said antigens to local immune cells. This surveillance feature of the epithelium is important for mounting appropriate immune responses to allergens and noxious substances (Iweala and Nagler, 2019). To more broadly understand how the CMA Donor 5 microbiota influences the gut mucosa, bulk RNA sequencing was performed on intestinal epithelial cells (IECs) of the ileum in GF mice, mice colonized with the CMA Donor 5 microbiota, and mice colonized with the microbiota of healthy infants (Donors 1-4) at one-week post-colonization. This data, which is a reanalysis of data from our previous study (Feehley et al., 2019) consists of an epithelial preparation that includes cells from the Peyer’s patches, as they are not excised in the isolation protocol we utilized (see Methods). Analysis of this data revealed hundreds of genes that were differentially expressed between CMA Donor 5-colonized mice and mice colonized with the healthy microbiota or left GF (**Table S1**). Many of these genes are involved in numerous immunological processes, including regulation of the epithelial barrier in response to bacterial colonization (*Defa17*, *Defa23*, *Il18*, *Il15*), antigen presenting cell (APC) function (*Cd8a*, *Cd83*), B cell responses (*Jchain*, *Fcgrt*) and type 2 responses (*Il13ra1*, *Il33*) (Harrison et al., 2015; Ganz et al., 2003; Nowarski et al., 2015, Campbell et al., 2021) (**Table S1**).

Additionally, several genes that have been previously described to be associated with a proinflammatory Th17 signature were found to be significantly upregulated in CMA Donor 5- colonized mice compared to healthy-colonized mice (Atarashi et al., 2015; Haberman et al., 2014). These genes include *Saa1*, *Saa3*, *Slc6a14*, *Fut2*, *Reg3b*, and *Reg3g* (**Figure 1B, Table S1**). *Saa1* is of particular interest because it is well known as an acute-phase protein expressed by hepatocytes during injury and infection (Sun and Ye, 2016). In the clinic, SAA-1 has been used as a biomarker of systemic inflammation for patients with Crohn’s disease as well as in patients with allergic asthma (Yarur et al., 2016; Bich et al., 2022; Ozseker et al., 2006). More recently, it has been shown be induced in the intestinal and bronchial epithelium and can regulate type 2 and type 3 inflammation at those sites (Lee et al., 2020; Smole et al, 2020). Furthermore, *Saa1* can be induced in the terminal ileal epithelium by Segmented Filamentous Bacteria (SFB), *Citrobacter rodentium*, *Escherichia coli* O157, and other epithelial-adherent bacteria (Atarashi et al., 2015).

Because *Saa1* has this documented affiliation with inflammatory responses and mice colonized with the CMA Donor 5 microbiota have inflammatory presentations, we hypothesized that *Saa1* induced by the Donor 5 microbiota may be a biomarker for a subset of patients with cow’s milk allergy. To determine if *Saa1* upregulation was induced by the microbiota of other CMA infants, GF mice were colonized with repository feces from CMA Donors 5-8, as well as repository feces from healthy Donors 1-4, and were euthanized at one-week post-colonization. Expression of *Saa1* was induced by colonization with all donor microbiotas compared to GF controls (**Figure 1C**). However, mice colonized with the CMA Donor 5 microbiota exhibited significantly higher expression of *Saa1* compared to all other CMA donors, all healthy donors, and GF mice. To determine the relevance of SAA-1 expresison to human infants, we quantified SAA- 1 in the serum, as well as antibodies specific for n Bos d 5, an epitope of BLG, in a larger cohort of 30 healthy and 30 CMA pediatric patients. As expected, CMA infants had significantly higher levels of antibodies against nBos d 5, a diagnostic indicator of cow’s milk allergy, in comparison to healthy infants. Interestingly, these infants also had significantly greater levels of serum SAA- 1 compared to healthy infants (**Figure 1D-E**). CMA Donor 5 exhibits a unique microbiota that can induce upregulation of *Saa1* in the ileal epithelium, which the microbiotas from other CMA infants failed to do. The CMA Donor 5 microbiota also induces a proinflammatory response in the ileum epithelium that includes the upregulation of Th2- and Th17-related genes.

### *Saa1* induction in the ileum epithelium of Donor 5-colonized mice is regulated by TLR4 signaling

After observing microbially induced upregulation of *Saa1* in the ileal epithelium of CMA Donor 5-colonized mice, we hypothesized that intestinal inflammatory responses mediated by SAA-1 might contribute to the occurrence of FA. Therefore, we next sought to understand the host-microbe interactions that mediate this phenotype. Analysis of fecal samples from healthy and CMA infant donors revealed that the microbial signature associated with healthy infants that were protected from FA consisted of enrichment of Clostridial species, particularly *Lachnospiraceae* (Feehley et al., 2019). In contrast, the microbial signature of CMA infants was highly enriched in Bacteroidetes. Interestingly, SAA1 has been previously correlated with increased abundance of *Bacteroides* species in a multi-omic analysis examining transcriptomic and microbial sequencing data from ileal biopsies taken from patients with inflammatory bowel disease (Tang et al., 2017). Furthermore, a previous study found Bacteroidetes to be enriched in two cohorts of infants with increased prevalence of allergy and autoimmunity compared to a genetically similar cohort with decreased prevalence of these diseases (Vatanen et al., 2016). Moreover, this study showed that lipopolysaccharide (LPS) from several Bacteroidetes species was hypoimmunogenic and was not protective against the onset of diabetes in a mouse model, as opposed to LPS from *Escherichia coli* which was hyperimmunogenic and protective.

SAA-1 has been shown to bind to LPS, opsonize Gram-negative bacteria and mediate their phagocytosis (Cheng et al., 2018; Shah et al., 2006). Furthermore, SAA-1 can induce inflammatory cytokine expression in macrophages in a TLR4-dependent manner (Niemi et al., 2011). TLR4, the host receptor that recognizes LPS, is known to regulate allergic inflammation in multiple models of atopic diseases through various mechanisms (Eisenbarth et al., 2002; Bashir et al., 2004; McAlees et al., 2015). Therefore, we looked at the role of TLR4 in inducing ileal expression of *Saa1*, our reproducible biomarker in CMA Donor 5-colonized mice. GF C57BL/6 mice with TLR4 sufficiency and deficiency were colonized at weaning with the CMA Donor 5 microbiota or the healthy Donor 1 microbiota and were euthanized one week later to examine *Saa1* expression in ileal IECs. Results showed that *Saa1* was significantly upregulated in mice colonized with the CMA Donor 5 microbiota compared to GF mice and mice colonized with the healthy Donor 1 microbiota when TLR4 was sufficient, but not when it was deficient (**Figure 2A)**. Along with *Saa1*, *Reg3b* has previously been associated with a Th17 signature induced by bacterial adherence to the ileal epithelium (Atarashi et al., 2015). Thus, we examined the expression of *Reg3b* in CMA Donor 5-colonized and healthy Donor 1-colonized mice as well. We observed that colonization of mice with the CMA Donor 5 microbiota and the healthy Donor 1 microbiota induced significant upregulation of *Reg3b* compared to GF mice (**Figure 2B)**. However, in the global absence of TLR4 signaling, mice colonized with the CMA Donor 5 microbiota did not have significant upregulation of *Reg3b* compared to GF mice. Interestingly, Donor 1-colonized mice still exhibited significant upregulation of *Reg3b* compared to GF and Donor 5-colonized mice. These results show that TLR4 signaling regulates the expression of *Saa1* and *Reg3b* induced by the CMA Donor 5 microbiota.

### Ileal regulatory T cells and Th17 cells are induced by the Donor 5 microbiota in a TLR4- dependent manner

Because SAA-1 has been previously implicated in regulating type 3 inflammation (Lee et al., 2020), the induction of *Saa1* in a TLR4-dependent manner would suggest that TLR4 also regulates immune cell populations. Therefore, we examined Foxp3^+^ regulatory T cells (Tregs), RORγt^+^ Foxp3^+^ Tregs, and IL-17A^+^ RORγt^+^ Th17 cells in TLR4-sufficient and -deficient mice (**Figure 2C-F**). Regulatory T cells are important for suppressing intestinal inflammation and are identified by the expression of the transcription factor Foxp3 (Bae et al., 2016). Foxp3^+^ Tregs that develop extrathymically are specifically induced by the microbiota and express the transcription factor RORγt (Sefik et al., 2015). RORγt^+^ Foxp3^+^ Tregs have been found to be especially important for the suppression of type 2 inflammation in several mouse models (Abdel-Gadir et al., 2019; Ohnmacht et al., 2015; Josefowicz et al., 2012). Thus, we examined the proportion and absolute numbers of Foxp3^+^ Tregs and RORγt^+^Foxp3^+^ Tregs in the ileum of mice colonized with the CMA and healthy microbiotas. GF C57BL/6 TLR4-sufficient and TLR4-deficient mice were colonized with the CMA Donor 5 microbiota or healthy Donor 2 microbiota at weaning. These mice were then euthanized one week later to isolate lymphocytes from the ileal lamina propria. Using the gating strategy shown in **Figure S2**, flow cytometric analysis revealed that, in TLR4-sufficient, but not TLR4-deficient, mice, both the frequency and numbers of Foxp3^+^ Tregs was significantly increased in CMA Donor 5-colonized mice compared to healthy Donor 2-colonized mice and GF mice (**Figure 2C-D).**

For RORγt^+^ Foxp3^+^ Tregs, there was an increase in frequency in CMA Donor 5-colonized mice compared to healthy Donor 2-colonized mice and GF mice when TLR4 signaling was sufficient (**Figure 2E**). Furthermore, there was a nearly significant decrease in the frequency of this population in TLR4-deficient mice colonized with the CMA microbiota. The absolute number of RORγt^+^Foxp3^+^ Tregs was significantly higher in TLR4-sufficient, CMA-colonized mice compared to GF mice of the same genotype (**Figure 2F**). There was also significant reduction in the absolute number of RORγt^+^Foxp3^+^ Tregs in TLR4-deficient, CMA-colonized mice compared to their wildtype counterparts. This population was also significantly reduced in healthy-colonized mice. Examining Th17 cell populations, both the frequency and number of Th17 cells were significantly increased in TLR4-sufficient, CMA-colonized mice compared to GF mice (**Figure 2G- H**). Importantly, the frequency and numbers of Th17 cells was significantly decreased in TLR4- deficient mice colonized with the CMA-associated microbiota compared to their TLR4-sufficient counterparts. Furthermore, in TLR4-deficient mice, there was a significantly higher frequency of Th17 cells in healthy-colonized and CMA-colonized mice compared to GF mice. The absolute number of Th17 cells was also significantly reduced in TLR4-deficient GF and healthy-colonized mice compared to their WT counterparts. Overall, this data shows that TLR4 regulates the induction of Th17 cells and Tregs in mice colonized with the microbiota from CMA Donor 5. In agreement with what has been observed in the literature, *Saa1* and *Reg3b* expression is coregulated with the induction of Th17 cells (Atarashi et al., 2015, Sano et al., 2015; Lee et al., 2020).

### TLR4 signaling in CD11c^+^ cells regulates the expression of Th17 genes, B cell genes, and alarmins ileum epithelium of Donor 5-colonized mice

The observation that TLR4 signaling not only regulates the induction of *Saa1* expression by the CMA Donor 5 microbiota but also Th17 and Treg subsets suggests that it may regulate intestinal inflammation that could exacerbate allergic responses to food. It has been well documented that TLR4 signaling in various cellular compartments differentially regulates allergic inflammation (Hammad et al., 2015; McAlees et al., 2015). In particular, CD11c^+^ DCs that are activated by SAA-1 promote IL-17A production in CD4^+^ T cells, contributing to Th2 and Th17 inflammatory responses in a model of allergic asthma (Ather et al. 2011). Furthermore, DCs have specifically been shown to regulate T and B cell responses through binding of retinol-SAA-1 complexes in the intestine (Bang et al., 2021). Thus, we examined the role of TLR4 signaling in CD11c^+^ cells in inducing *Saa1* and *Reg3b*, as well as induction of Tregs and Th17 cells. To do this, we used mice with a conditional knockout of TLR4 in CD11c^+^ cells (CD11c^cre^TLR4^fl/fl^). Because these mice were housed under specific-pathogen free (SPF) conditions, we first depleted their microbiota by treating pups with a cocktail of antibiotics (see Methods) by intragastric gavage for seven days prior to weaning. At weaning, these pups were colonized with the CMA Donor 5 microbiota or the healthy Donor 1 microbiota or left uncolonized as SPF controls. Mice were then sacrificed one week later to examine gene expression in ileal IECs. All experiments were littermate controlled.

In mice with a conditional mutation of TLR4 in CD11c^+^ cells, we observed changes in ileal gene expression that were similar to those seen in global TLR4 knockout mice. In CD11c^WT^ mice, both *Saa1* and *Reg3b* were significantly upregulated in the ileal epithelium of CMA Donor 5- colonized mice compared to healthy Donor 1-colonized mice and SPF controls (**Figure 3 A-B**). However, in their CD11c^ΔTLR4^ littermates, this upregulation was no longer observed. Because CD11c^+^ cells have been shown to regulate intestinal IgA through SAA-1 (Bang et al., 2021), we also examined the expression of the heavy chain of IgA (*Igha*) and *Jchain*. We observed that *Igha* and *Jchain* were both significantly upregulated in the ileal epithelium of CMA Donor 5-colonized, CD11c^WT^ mice but not in their CD11c^ΔTLR4^ littermates (**Figure 3C-D**, note that our Method for isolation of IECs does not include removal of Peyer’s patches). These observations correlated with a significant reduction in fecal IgA in CD11c^ΔTLR4^ mice compared to CD11c^WT^ mice colonized with the CMA-microbiota (**Figure 3E**).

**Figure 3:**
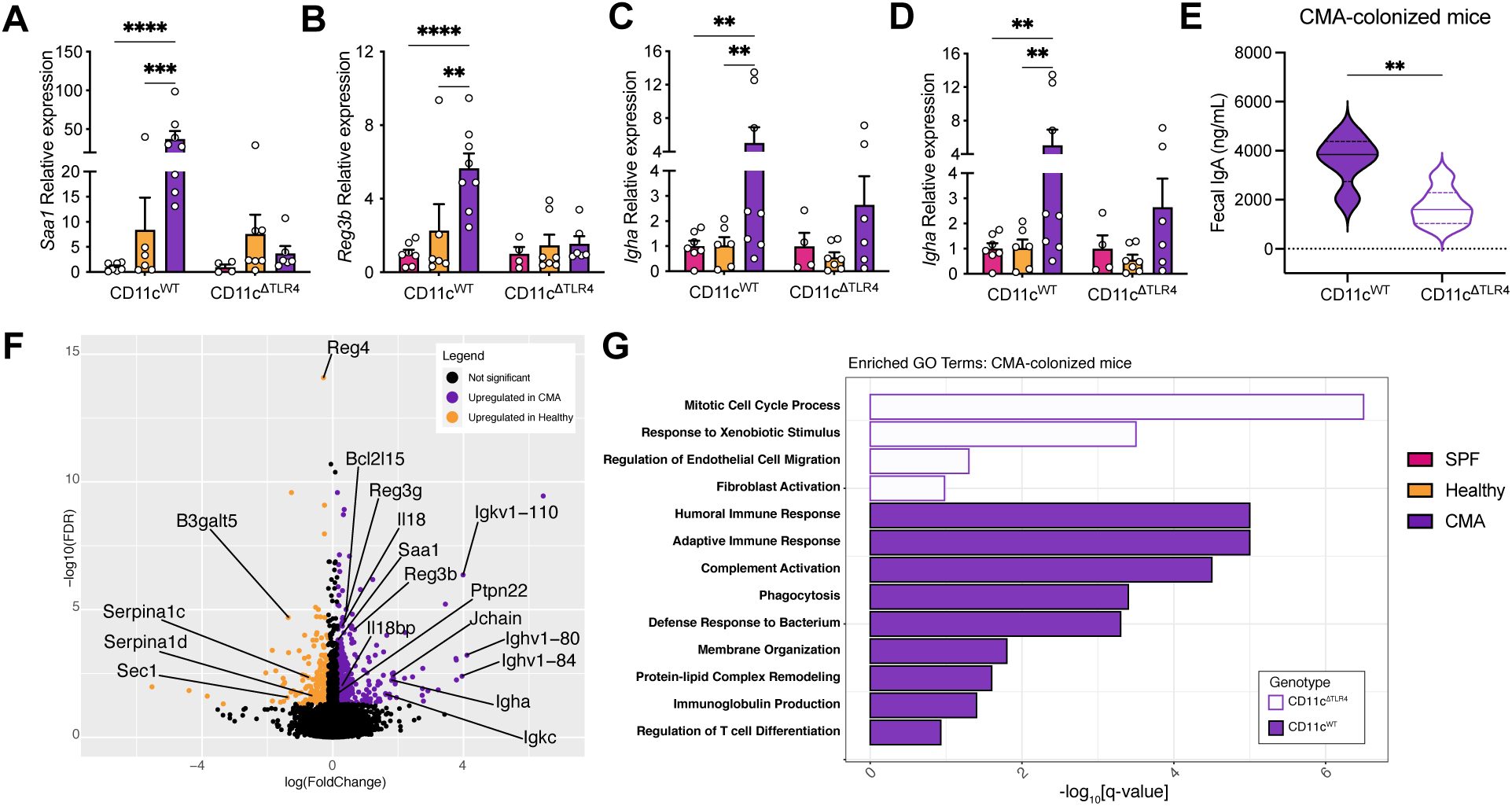
TLR4 signaling in the CD11c compartment regulates ileal Th17-associated genes and B cell genes in CMA-colonized mice in an antibiotic-treatment model Relative expression of A) *Saa1* B) *Reg3b*, C) *Igha* and D) *Jchain* in antibiotic-treated SPF mice colonized with the CMA microbiota, healthy microbiota, or left uncolonized after treatment one-week post-colonization/weaning. E) Fecal IgA levels in antibiotic-treated, CMA-colonized CD11c^WT^ and CD11c^Δ*Tlr4*^ littermates F) Volcano plot of significantly upregulated genes in antibiotic-treated SPF mice colonized with the CMA or healthy microbiotas. G) Identified gene ontology (GO) pathways significantly enriched in ileal IECs of CMA-colonized CD11c^WT^ and CD11c ^Δ*Tlr4*^ mice. Data are pooled from at least two independent experiments per condition. Statistics were calculated using 2-way ANOVA with multiple testing using the Benjamini, Krieger, and Yekutili method. \*\**P* < 0.01, \*\*\**P* < 0.001, \*\*\*\**P* < 0.0001

In our previous RNA sequencing data, *Il33* was found to be significantly upregulated in the ileal epithelium of CMA Donor 5-colonized mice (**Table S1**). Therefore, we examined the expression of the classical epithelial alarmins *Il33*, *Il25*, and *Tslp*, which play a role in mediating type 2 inflammation in response to damage and sensitization with an allergen by qRT-PCR (Peterson et al., 2014). In CD11c^WT^ mice, *Il33* was nearly significantly upregulated in CMA- colonized mice compared to both healthy-colonized mice and SPF mice, which was not seen in CD11c^ΔTLR4^ littermates (**Figure S3A**). Conversely, *Il25* was not observed to be upregulated in CD11c^WT^ mice colonized with the CMA microbiota (**Figure S3B**). However, in CD11c^ΔTLR4^ mice, *Il25* expression was significantly upregulated in the ileal epithelium of CMA-colonized mice compared to both healthy-colonized mice and SPF controls. The expression of *Tslp* was not upregulated in any of the mice examined (data not shown). These results suggest that TLR4 signaling in CD11c^+^ cells may regulate type 3 responses, type 2 responses, and B cell responses in the ileum of CMA Donor 5-colonized mice and validates results obtained from the bulk RNA sequencing data performed in ex-GF mice (**Tables S1**).

To better understand how TLR4 signaling in CD11c^+^ cells regulates gene expression in the ileal epithelium of CMA Donor 5-colonized mice, bulk RNA sequencing was conducted on RNA isolated from ileal IECs of antibiotic-treated CD11c^WT^ and CD11c^ΔTLR4^ mice that were uncolonized SPF controls, colonized with the CMA microbiota, or colonized with the healthy microbiota. Differential expression analysis between CD11c^WT^ mice colonized with the CMA microbiota and the healthy microbiota revealed several genes to be specifically upregulated in each group. Some of the genes upregulated in this dataset were also upregulated in our previous RNA sequencing dataset which analyzed ileal IEC gene expression in ex-GF mice colonized with each of the four healthy infant donors compared to CMA Donor 5 (see **Tables S1 and S2**). Specifically, the expression of *Saa1* was upregulated, along with other previously identified Th17-associated genes, such as *Reg3b* and *Reg3g*, (**Figure 3F**, **Table S2**). Other genes involved in epithelial-induced proinflammatory responses and barrier function such as *Il18* and *Il18bp* were also upregulated. Many genes related to B cell responses were upregulated, including *Igha*, *Jchain*, and numerous immunoglobulin genes.

Next, pathway analysis was conducted on CD11c^WT^ and CD11c^ΔTLR4^ mice colonized with the CMA Donor 5 microbiota. Results showed that many genes upregulated in CD11c^WT^ mice were involved in immunological processes. In agreement with the differential expression analysis and qRT-PCR data, genes involved in humoral immune responses and immunoglobulin production were highly significantly upregulated (**Figure 3G**). Genes involved in regulation of T cell differentiation were also significantly upregulated, in agreement with our previous data showing that global TLR4 signaling regulates several T cell subsets in CMA Donor 5-colonized mice (**Figures 2C-H**). Interestingly, genes involved in defense against bacteria were also upregulated, suggesting that members of the CMA Donor 5 microbiota may be antigenically proinflammatory in nature. In CD11c^ΔTLR4^ mice, these immunological pathways were not upregulated. Instead, the pathways upregulated were more aligned with homeostatic regulation of the epithelium.

To better understand the bacterial communities that induce these genes in CMA- colonized and healthy-colonized mice, 16S sequencing was performed on samples of ileal contents acquired from these mice at euthanasia. We are aware that because these mice originated from a SPF colony and were treated with antibiotics, the gut microbiota will consist of an amalgamation of bacteria originating from the SPF microbiota and human-derived bacteria from the infant donors. Therefore, we first performed a basic, holistic analysis of alpha and beta diversity to assess the ecological dynamics of the commensal microbiota and determine the fidelity of transfer of the human infant microbiota. The Shannon index was used to calculate diversity within each sample. Analysis of each sample by group determined that the average diversity within each sample was not significantly different between colonization status (**Figure S3C**). To determine the fidelity of transfer, beta diversity was determined using the Bray-Curtis index of dissimilarity. Analysis between groups showed that there were significant differences between microbial communities based on colonization status (**Figure S3D**). This suggests that the transfer of the human infant gut microbiota was successful following antibiotic treatment of SPF mice and that differences in gene expression observed in CMA-colonized mice and healthy- colonized mice were due to the infant microbiota. In summary, these data show that in our antibiotic-treated SPF model, the phenotypic upregulation of *Saa1* and *Reg3b* (as well as numerous other immune system genes) in isolated ileal IECs and the production of fecal IgA, is most likely due to the engraftment of the CMA Donor 5 microbiota.

### Th17- and B cell-associated genes are regulated by TLR4 signaling in CD11c^+^ cells in ex-GF mice colonized with the CMA microbiota

We have shown that global and conditional expression of TLR4 in CD11c^+^ cells regulates ileal IEC gene expression in CMA-colonized mice in both GF and antibiotic-treated colonization models, respectively. In order get an understanding of members of the CMA Donor 5 microbiota that may contribute to these immune responses, we proceeded to rederive SPF CD11c^WT^ and CD11c^ΔTLR4^ mice as GF. The experiments that were performed in the antibiotic-treated model were repeated using these GF mice following the same experimental protocol utilized in the global TLR4 knockout mice. The mice were colonized at weaning with the CMA Donor 5 or healthy Donor 2 microbiota and euthanized one-week post-colonization to isolate RNA from ileal IECs and examine *Saa1* expression.

Expression analysis showed results that were strikingly similar to those observed in SPF global and CD11c-conditional TLR4 knockout mice. *Saa1* and *Reg3b* were significantly upregulated in CD11c^WT^ CMA-colonized mice compared to healthy-colonized and GF mice (**Figure 4A-B**). However, in CMA-colonized mice with a conditional mutation in TLR4 in CD11c^+^ cells, these genes were no longer upregulated. When looking at B cell-related genes, we similarly observed that the expression of *Igha* and *Jchain* was higher in CMA-colonized mice compared to healthy- colonized and GF mice, which was regulated by TLR4 signaling in CD11c^+^ cells (**Figure 4C-D**). This translated to differences in the production of fecal IgA. In CD11c^WT^ mice, fecal IgA was significantly higher in CMA-colonized mice compared to healthy-colonized and GF mice (**Figure 4E**). In CD11c^ΔTLR4^ mice, there was significantly higher fecal IgA in CMA-colonized and healthy- colonized mice compared to GF mice. Interestingly, there was a significant reduction in fecal IgA in CMA-colonized CD11c^ΔTLR4^ mice compared to their CD11c^WT^ littermates. This significant reduction was not observed between CD11c^WT^ and CD11c^TLR4^ mice colonized with the healthy microbiota or CD11c^WT^ and CD11c^ΔTLR4^ GF mice. This data shows that TLR4 signaling in CD11c^+^ cells regulate the expression of *Saa1* and *Reg3b* in ileal IECs of CMA Donor 5-colonized mice in both the GF and antibiotic-treated colonization models. Furthermore, in CMA Donor 5-colonized mice, TLR4 signaling in CD11c^+^ cells regulate B cell responses, and thus intestinal IgA production.

**Figure 4:**
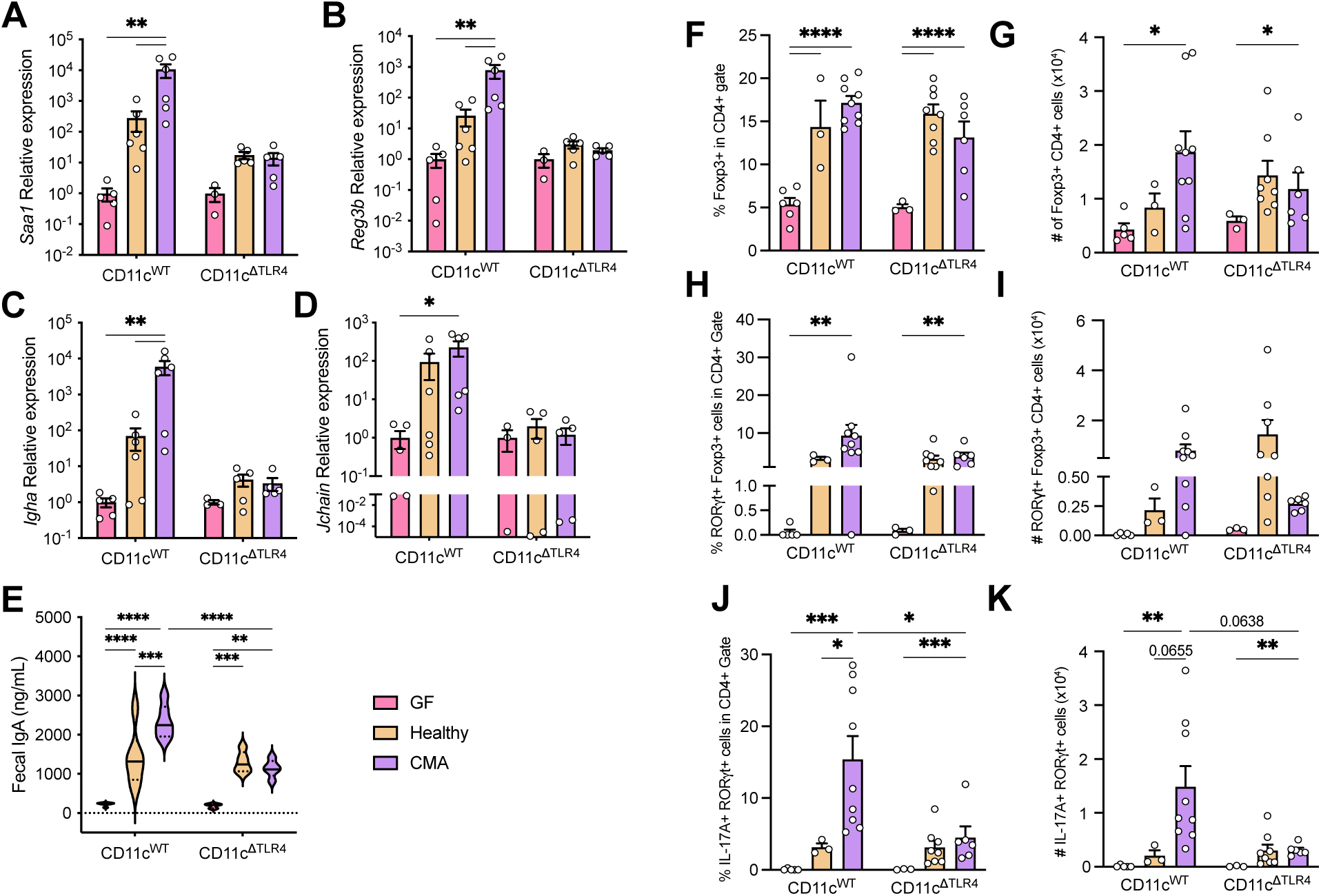
Conditional KO of TLR4 in the CD11c compartment regulates ileal gene expression, Th17 cells, and fecal IgA in CMA-colonized mice in an ex-GF model Relative expression of A) *Saa1*, B) *Reg3b*, C) *Igha*, and D) *Jchain* in GF, CMA-colonized, and healthy colonized mice one-week post-colonization. E) Fecal IgA levels of GF, healthy-colonized, and CMA-colonized CD11c^WT^ and CD11c^Δ*Tlr4*^ mice. Frequency and absolute numbers of F-G) Foxp3^+^ Tregs, H-I) Foxp3^+^ RORγt^+^ Tregs, and J-K) IL-17^+^ RORγt^+^ Th17 cells in the ileal tissue of GF, healthy-colonized, and CMA-colonized CD11c^WT^ and CD11c^Δ*Tlr4*^ mice one- week post-colonization/weaning. Data are pooled from at least two independent experiments per condition. Statistics were calculated using 2-way ANOVA with multiple testing using the Benjamini, Krieger, and Yekutili method. \**P* < 0.05, \*\**P* < 0.01, \*\*\**P* < 0.001, \*\*\*\**P* < 0.0001

### TLR4 signaling in CD11c^+^ cells regulates Th17 populations but not Treg populations in the ileum of CMA Donor 5-colonized mice

Because similar gene expression results were seen in ex-GF mice with TLR4 globally knocked out, we proceeded to examine whether TLR4 signaling in CD11c^+^ cells also regulated T cell populations. We observed that TLR4 signaling in CD11c^+^ cells did not regulate T cell populations in the same way that global TLR4 signaling did. Both CMA and healthy-colonized mice had significantly more frequent Foxp3^+^ Tregs compared to GF mice when TLR4 signaling was sufficient in CD11c^+^ cells (**Figure 4F**). The absolute number of Foxp3^+^ Tregs was also significantly higher in CD11c^WT^, CMA-colonized mice compared to their GF counterparts (**Figure 4H**). The frequency of allergy suppressive RORγt^+^ Foxp3^+^ Tregs was increased in CD11c^WT^ mice colonized with the CMA microbiota compared to GF mice (**Figure 4H**). However, the frequency and absolute number of both Foxp3^+^ Tregs and RORγt^+^Foxp3^+^ Tregs was not different between CD11c^WT^ and CD11c^ΔTLR4^ littermates in all colonization conditions (**Figure 4F-I**). This suggests that TLR4 signaling in another cellular compartment regulates the Treg populations that are induced by the CMA Donor 5 microbiota.

Examining Th17 populations, we observed that in CD11c^WT^ mice, there were significantly higher frequencies in healthy and CMA-colonized mice compared to GF mice (**Figure 4J**). Furthermore, the frequency of this population was significantly reduced in CD11c^ΔTLR4^ mice colonized with the CMA microbiota compared to their wildtype littermates. This reduction was also observed in the absolute numbers of Th17 cells (**Figure 4K**). The absolute number of Th17 cells in CMA-colonized mice was also greater compared to healthy-colonized and GF mice when TLR4 signaling was sufficient in CD11c^+^ cells. In CD11c^ΔTLR4^ mice, the frequency and absolute number of Th17 cells was significantly higher in CMA-colonized mice compared to GF mice (**Figure 4K**). These results show that in CMA-colonized mice, TLR4 signaling in CD11c^+^ cells regulates Th17 but not Treg populations. This agrees with previous findings that SAA-1 induces Th17 differentiation only in the presence of CD11c^+^ DCs from the lamina propria (Ivanov et al., 2009). Together, these data show that while global TLR4 signaling regulates the induction of Tregs and Th17 cells in CMA Donor 5-colonized mice, TLR4 signaling in CD11c^+^ cells specifically regulates the induction of Th17 cells.

### CMA Donor 5-colonized mice have higher representation of Bacteroidetes in their ileal contents

Rederivation of GF mice with TLR4 conditionally knocked out in CD11c^+^ cells allowed us to examine microbes that may contribute to *Saa1* expression, an analysis which was not possible in our previously utilized antibiotic-treatment model. To determine microbial contributors to the observed TLR4-dependent phenotypes, we conducted 16S sequencing on ileal samples of CMA and healthy-colonized CD11c^WT^ and CD11c^ΔTLR4^ mice. Through analysis of these sequences, we determined that multiple taxa were differentially abundant between healthy and CMA-colonized mice. In healthy-colonized mice several Clostridial taxa were significantly more abundant, including *Lachnospiraceae* and *Clostridium Cluster XIVa*, which we have previously reported to be barrier protective Clostridia (Stefka et al., 2014; Feehley et al., 2019) (**Figure S4A**). Importantly, all Gram-negative bacteria that were identified as differentially abundant were significantly higher in CMA-colonized mice compared to healthy-colonized mice (**Figure S4B**).

After obtaining these results, we followed up with shotgun metagenomic sequencing of DNA from the same samples to get a more resolved understanding of which species were differentially abundant between healthy Donor 2-colonized and CMA-Donor 5 colonized mice. Taxonomic analysis showed that the bacterial communities in healthy Donor 2-colonized, and CMA Donor 5-colonized mice were vastly different (**Figure 5A**). In agreement with the 16S data, CMA Donor 5-colonized mice had an expansion of Gram-negative bacteria, such as Proteobacteria and Bacteroidetes. Healthy-colonized mice had a higher abundance of *Enterococcus* and *Lachnospiraceae*, which was also in agreement with the 16S data. As expected, colonization of GF mice with the CMA or healthy microbiota was observed to be distinct according to beta-diversity metrics (**Figure 5B**).

**Figure 5:**
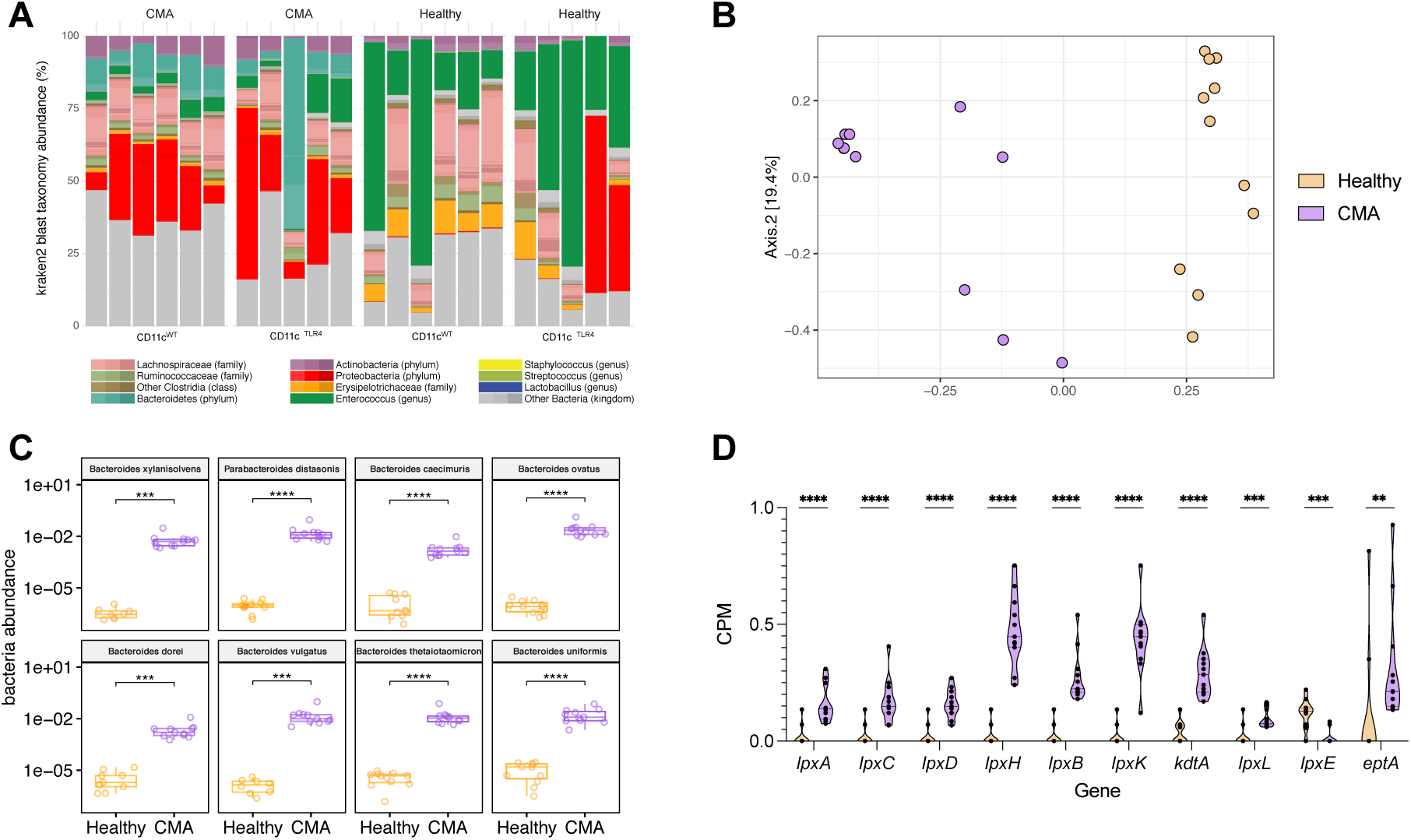
CMA-colonized mice harbor an increased abundance of *Bacteroidetes* and other Gram-negative bacteria in ileal contents. A) Relative abundance of metagenomic reads as classified by Kraken2 in healthy- and CMA- colonized mice with and without conditional knockout of TLR4 in CD11c^+^ cells at one-week post-colonization B) Beta diversity of healthy Donor 2-colonized and CMA Donor 5-colonized mice using the Bray-Curtis metric. C) Differentially abundant Bacteroidetes species between CMA- and healthy-colonized mice. D) Counts per million of genes involved in KEGG pathways related to KDO2-lipid A biosynthesis and modification. Statistics were calculated by performing Wilcoxon rank sum test between both conditions.

Differential abundance analysis of shotgun metagenomic sequences revealed numerous bacterial species to be particularly enriched in healthy Donor 2-colonized or CMA Donor 5- colonized mice. Interestingly, we again saw that all Gram-negative species were significantly more abundant in CMA Donor 5-colonized mice than healthy Donor 2-colonized mice, which agreed with the 16S sequencing data we previously collected (**Figure 5C, Figure S5**). Specifically, CMA Donor 5-colonized mice had an increase in Bacteroidetes compared to healthy-colonized mice (**Figure 5C**). The Bacteroidetes are a phylum of bacteria that have previously been associated with allergy (Vatanen et al., 2016). Bacteroidetes produce hypoimmunogenic LPS compared to LPS from Escherichia coli, which may contribute to allergies and autoimmunity through lack of differential development of endotoxin tolerance (Martin et al., 2001). Knowing this, we analyzed the representation of LPS synthesis and LPS modification genes in these samples. We saw that all genes analyzed were significantly more represented in CMA-colonized mice, with exception to lpxE, which was significantly higher in healthy-colonized mice (**Figure 5D**). Together, these data show that the Donor 5 microbiota has higher abundance of Gram-negative bacteria, particularly Bacteroidetes, in comparison to the Donor 2 microbiota. Accordingly, the CMA Donor 5 microbiota also has increased representation of LPS genes that are likely derived from Bacteroidetes.

### TLR4 regulates type 2 and type 3 T cells and ILCs during early development of an allergic response in CMA-colonized mice

Cumulatively, the CMA Donor 5 microbiota induces immunological responses in the ileum that are proinflammatory in nature. In response to the CMA Donor 5 microbiota, compensatory defense mechanisms are enacted through the induction of IgA (**Figure 3, 4**) and antimicrobial peptides (*Reg3b*, **Figure 4**) and these responses are regulated by TLR4 in CD11c^+^ cells. Since these responses are induced by a microbiota originating from an infant with a severe allergy to cow’s milk, one question that remains is the role of this microbiota in inducing allergic inflammation through TLR4-dependent mechanisms. To investigate this, we sensitized TLR4-sufficient and TLR4-deficient mice colonized with the CMA Donor 5 microbiota, the healthy Donor 2 microbiota, or GF controls. Using a model where mice are sensitized twice with BLG plus the mucosal adjuvant cholera toxin (CT), we looked at ileal ILC and T helper cell populations in the early development of an allergic response. Specifically, using the gating strategy presented in **Figure S6A**, we examined ILC2 and Th2 cells to examine mediators of type 2 inflammatory responses, as well as ILC3s and Th17 cells to examine mediators of type 3 inflammation. Following this sensitization protocol, we observed significant increases in total ILCs and T cells in healthy-colonized mice compared to GF and CMA-colonized mice when TLR4 signaling was deficient, as well as their WT counterparts (**Figure S6B-E**).

Furthermore, in TLR4 sufficient mice, we observed GF and CMA Donor 5-colonized mice had a greater frequency of ILC2s compared to healthy colonized mice (**Figure 6A**). This was also observed in TLR4-deficient mice. This agrees with previous findings that both GF and CMA- colonized mice are more susceptible to allergic sensitization than healthy-colonized mice (Feehley et al., 2019). Interestingly, the frequency of ILC2s in both GF and CMA-colonized mice was significantly increased in TLR4-deficient mice compared to TLR4-sufficient mice while the frequency in healthy Donor 2-colonized mice did not increase. The absolute number of ILC2s increased in all colonization statuses when TLR4 was deficient compared to their WT counterparts (**Figure 6B**). Additionally, Th2 cells were greatly induced in all colonization conditions when TLR4 signaling was deficient but not when it was sufficient (**Figure S6F-G**), which agrees with previous observations in TLR4-deficient mice (Bashir et al., 2004).

**Figure 6:**
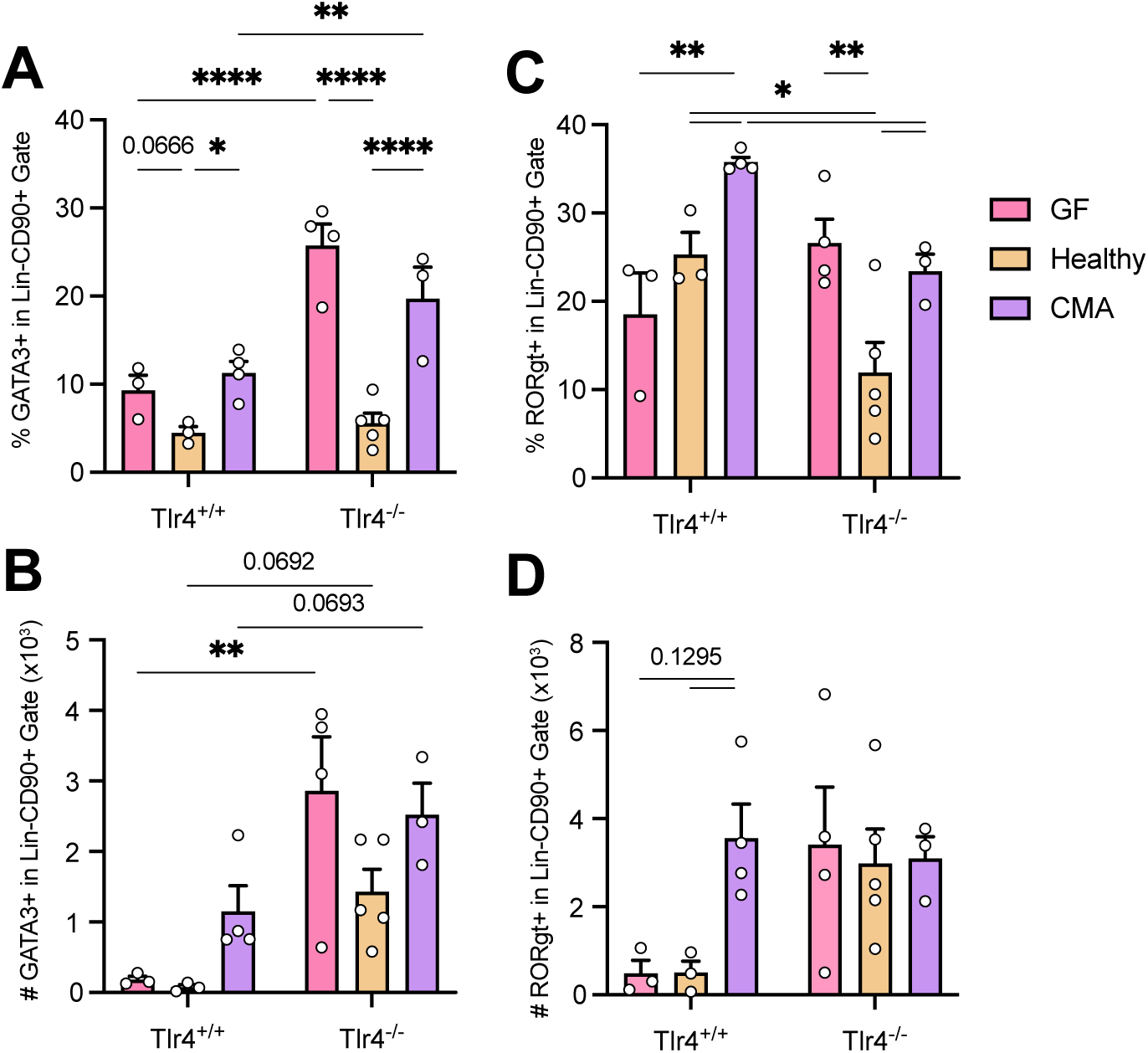
The CMA microbiota induces innate type 2/type 3 inflammation in the early development of an allergic response. A, C) Frequency and B, D) absolute numbers of A-B) ILC2s and C-D) ILC3s in the ileum lamina propria of GF, healthy Donor 2-colonized, and CMA Donor 5-colonized *Tlr4*^+/+^ and *Tlr4*^-/-^ mice. Pooled data from at least 2 independent experiments per condition. Statistics were calculated using 2-way ANOVA with multiple testing using the Benjamini, Krieger, and Yekutili method. \**P* < 0.05, \*\**P* < 0.01, \*\*\*\**P* < 0.0001

When examining ILC3s, we observed a significant increase in frequency of ILC3s in CMA- colonized mice compared to healthy-colonized and GF mice when TLR4 signaling was sufficient (**Figure 6C**). Furthermore, this frequency was significantly decreased when TLR4 was deficient. Similar to what was seen in the ILC2s, the frequency of ILC3s was significantly higher in GF and CMA-colonized mice compared to healthy-colonized mice when TLR4 was deficient. The absolute numbers of ILC3s were not different between genotypes and colonization status, although CMA- colonized mice did have an increase in numbers (**Figure 6D**). The frequency of Th17 cells was increased in TLR4-deficient GF and CMA-colonized mice compared their WT counterparts, while no differences were seen in the absolute numbers (**Figure S6H-I**). Together, this data shows that TLR4 signaling regulates the frequencies of ILC3s and Th17 cells in GF and CMA-colonized mice. In healthy-colonized mice, TLR4 signaling regulates ILC3 frequencies, but not Th17 frequencies.

In summary, early innate effectors (ILC2s and ILC3s) are induced in allergy susceptible CMA-colonized mice and GF mice. Following sensitization, ILC2s are induced similarly in GF and CMA-colonized mice, while ILC3s are more highly induced in CMA-colonized mice. In the absence of TLR4 signaling, ILC2s are increased in both GF and CMA-colonized mice, while ILC3s are greatly reduced only in CMA-colonized mice, suggesting that innate type 3 inflammation is regulated by TLR4 in these mice. When TLR4 signaling is sufficient, healthy-colonized mice have reduced ILC2s and ILC3s compared to CMA-colonized mice. In agreement with previous findings (Bashir et al., 2004), when TLR4 signaling is deficient, all colonization statuses experience an increase in frequency of Th2 cells. Moreover, in allergy susceptible GF and CMA-colonized mice, there is an increased frequency of Th17 cells that is not observed in healthy-colonized mice. These data show that TLR4 regulates a complex network of type 2 and type 3 inflammatory cells of the innate and adaptive arms.

## Discussion

In the data presented herein, we employ multiple colonization models to show that the CMA-associated microbiota induces a mixed inflammatory response in the ileum. This not only includes the induction of *Saa1* and *Reg3b*, but other inflammatory and mucosal genes, including *Il33*, *Il13ra, Il15*, *Il18* (Table S1). Furthermore, as a part of this inflammatory response, increased Tregs and Th17 cells are induced, along with fecal IgA, in comparison to healthy-colonized mice. Prior to epithelial cell activation elicited by sensitization with a dietary antigen, the CMA microbiota orients the ileal mucosa toward a type 2/type 3 inflammatory response. Adding insult to injury, it is unsurprising that upon cow’s milk consumption, CMA Donor 5 experienced severe symptoms with an intestinal mucosa that is already poised toward inflammation. The atopic symptoms that CMA Donor 5 suffered from within minutes after feeding (i.e., vomiting, urticaria), as well as the gastrointestinal inflammation (nodular hyperplasia and eosinophilic infiltration) are likely the outcome of a vigorous allergen-specific response that has developed within the first months of the infant’s life. In our study, we examined an early snapshot of the inflammatory responses that this microbiota induces.

In addition to the inflammatory transcriptional profile induced in the ileal mucosa, significantly elevated proportions, and numbers of Foxp3^+^ Tregs, RORγt^+^ Foxp3^+^ Tregs, and IL- 17A^+^ RORγt^+^ Th17 cells, as well as increased induction of IgA, were specific to mice colonized with the CMA Donor 5 microbiota and were regulated by TLR4 signaling. *Saa1* is induced in the ileal epithelium by epithelial adherent bacteria and is downstream of the IL-23 signaling axis initiated by intestinal DCs in response to microbial stimulation (Sano et al., 2015). Induction of *Saa1* by the epithelium increases the production of IgA by B cells and IL-17A in Th17 cells and leads to the exacerbation of intestinal inflammation caused by this T cell subset (Atarashi et al., 2015; Lee et al., 2020; Bang et al., 2021). Concordantly, in our studies, the induction of effector Th17 cells and IgA production by the CMA microbiota was coregulated with the induction of *Saa1* and dependent on TLR4 signaling in CD11c^+^ cells. Interestingly, Tregs were also highly induced in Donor 5-colonized mice. This is in alignment with previous studies observing robust induction of Foxp3^+^ Tregs and RORγt^+^ Foxp3^+^ Tregs in response to gut inflammation induced by Gram-negative bacteria, which required TLR4 signaling (Liu et al., 2022; Jia et al., 2017). The observation that the induction of Tregs is independent of TLR4 signaling in CD11c^+^ cells, also agrees with previous reports showing that induction of *Saa1* induces Th17 cells and intestinal IgA but not Tregs (Bang et al., 2021; Ivanov et al., 2009). With the production of SAA-1 favoring induction of Th17 cells, coupled with type 2 and type 3 inflammatory signals from the epithelium, the increased induction of Tregs is likely a compensatory mechanism to address inflammation within the local microenvironment.

Rederivation of CD11c^cre^TLR4^fl/fl^ mice also allowed us to identify bacteria enriched in CMA- colonized mice without contamination from the SPF commensal microbiota. Through use of this gnotobiotic model, we were able to determine that Gram-negative bacteria, particularly Bacteroidetes, are specifically enriched in the CMA microbiota. The Bacteroidetes are known to be common commensals of the mammalian intestine (Donaldson et al., 2016). While some have been documented to promote intestinal health (Mazmanian et al., 2008), others have been associated with intestinal inflammation, and particularly, the induction of *Saa1* in the ileal epithelium (Mills et al., 2022; Atarashi et al, 2015). Moreover, increased SAA-1 expression and Th17 cells have been correlated with increased abundance of *Bacteroides* in ileal biopsies from IBD patients (Tang et al., 2017). As *Saa1* has been shown to contribute to allergic inflammation (Smole et al., 2020), it is reasonable to suspect that induction of *Saa1* by a Bacteroidetes-enriched microbiota contributes to food allergies. While this study does not show induction of *Saa1* by Bacteroidetes, future investigations can be done to identify specific Bacteroidetes taxa from the CMA microbiota that can induce *Saa1* in the epithelium.

Upon sensitization with BLG plus CT, ILC and T cell populations were differentially regulated in GF, healthy-colonized, and CMA-colonized mice. TLR4-sufficient healthy-colonized mice exhibit reduced ILC2s compared to GF and CMA-colonized mice, as well as an increased frequency of total T cells (**Figure 6A, S6E**). This is in alignment with the protective effects of the healthy microbiota (Feehley et al., 2019). On the other hand, CMA-colonized and GF WT mice, which are more susceptible to anaphylaxis than healthy mice, exhibited similar frequencies of ILC2s (**Figure 6A**). CMA-colonized mice had a higher number of ILC2s compared to GF mice which agrees with the observation that CMA-colonized mice have increased total ILCs (**Figure 6B, S6C**). During these early responses, Th2 immunity has not fully developed in WT mice, as dictated by the low frequency and abundance of Th2 cells in all colonization conditions (**Figure S6F-G**). Thus, during early allergic responses, both CMA-colonized and GF mice can initiate innate type 2 inflammation. However, unlike GF mice, CMA-colonized mice have increased induction of ILC3s (**Figure 6C**). While ILC3s are important for maintaining intestinal homeostasis through myriad mechanisms (Zhou and Sonnenburg, 2019), they can act as potent inducers of epithelial SAA-1 and type 3 inflammation through IL-22-dependent mechanisms (Sano et al., 2015; Gunasekera et al., 2020; Eken et al., 2014). The specific induction of both ILC2s and ILC3s would suggest that CMA-colonized mice have a mixed type 2/type 3 intestinal inflammatory response that does not occur in GF mice upon sensitization. In experimental mouse models, this mixed type 2/type 3 inflammatory response exacerbates antigen-specific allergic inflammation (Nakajima et al., 2014; He et al., 2007). Therefore, our data suggests that while both CMA-colonized and GF mice are both susceptible to allergic sensitization, the mechanisms by which they are susceptible are not the same. While GF mice are prone to atopy due to the dysregulation of IgE (Cahenzli et al., 2013; Hong et al., 2019), the CMA microbiota actively promotes inflammatory responses.

Abrogation of TLR4 signaling can increase allergic susceptibility (Bashir et al., 2004; Brandt et al., 2013). When TLR4 signaling is deficient, GF and CMA-colonized mice have increases in ILC2s compared to when TLR4 was sufficient (**Figure 6A-B**), confirming that the inability to signal through TLR4 further increases susceptibility to allergic sensitization Furthermore, TLR4-deficient mice of all colonization statuses have increased frequency of Th2 cells compared to their WT counterparts (**Figure S6F**). Unexpectedly, in comparison to their WT counterparts, healthy- colonized mice also have increased number of Th2 cells and ILC2s (**Figure 6B, S6G**). However, coupled with sensitization, this may be due to healthy-colonized mice also having a TLR4- dependent increase in total ILCs and T cells, which is not observed in GF and CMA-colonized mice (**Figure S6C, E)**. Interestingly, CMA colonized mice have decreased ILC3s, but increased Th17 cells compared to their WT counterparts (**Figure 6C, S6H**). This too may be an effect that is specific for the CMA microbiota coupled with sensitization. In response to epithelial adherent bacteria, Treg-specific deletion of MyD88 signaling, a downstream adaptor of TLR4, reduces intestinal IgA production, increases barrier permeability, and dysregulates the balance between Tregs and Th17 cells, leading to increased Th17 induction (Wang et al., 2015). Global abrogation of TLR4 could thus disrupt the immunological balance in the intestine through microbiota-dependent and -independent mechanisms. This suggests that TLR4 signaling is regulating the balance between the innate and adaptive arms during the early phases of allergic sensitization.

As very few studies show a proactive role of a microbiota in mediating inflammatory responses to food antigen, this work is novel in its demonstration of microbially-mediated induction of type 2/type 3 inflammation prior to and following sensitization with BLG. The inflammation that accompanies food allergies is multifaceted, and thus, defining etiological determinants poses as a notable difficulty in the study of the disease. Through this work, we characterize an intestinal inflammatory response that is the direct result of host-microbe interactions with the CMA Donor 5 microbiota. While Donor 5 exhibited an extreme allergic phenotype that did not occur in Donors 6-8, the observation that SAA-1 is elevated in the serum of a subset of infants in the larger cohort (**Figure 1E**) suggests that the intestinal inflammation that is associated with Donor 5 may be relevant to other CMA infants. Since FAs and other atopic diseases elicit diverse immunological responses, the use of SAA-1 as a biomarker could characterize a specific type of inflammatory response that is microbially mediated, poises the intestine toward type 2/type 3 immunity, and is rapidly intensified in an antigen-specific manner upon exposure to cow’s milk. This manuscript defines the features of an early immune response induced by an atopic microbiota that will be important to examine in subsequent food antigen-specific responses. Future work can examine the adaptive immunological responses that are regulated by TLR4 globally and specifically in CD11c^+^ cells in the induction of allergic sensitization to food antigen using a full sensitization model. Such experiments would be important for understanding the root of inflammatory manifestations that could be present in CMA infants with high serum SAA-1 and elucidate the cellular contributors of these inflammatory responses. In time, this could potentially lead to the utilization of more personalized clinical treatments for CMA patients with this type of allergic inflammation.

## Acknowledgments

We thank the staff of The University of Chicago Animal Resource Center, particularly the Gnotobiotic Research Animal Facility (GRAF) for their expert technical assistance. We would also like to like to thank the Duchossois Family Institute (DFI) for their assistance in bioinformatic analysis.

## Author Contributions

EC and CRN designed the study. EC performed mouse experiments with help from RTP, LMM, LAH, and AMK. EC and CRN analyzed results with assistance from the DFI. RBC cared for patients and provided patient samples. R.N. and L.P. collected and analyzed patient samples in Fig. 1. EC and CRN wrote the manuscript. All authors read and commented on the manuscript.

## Data Availability

The 16S rRNA-targeted and metagenomic sequencing raw FastQ data files have been deposited in the NCBI Sequence Read Archive (accession #SUB11826475). RNA sequencing raw FastQ data files have been deposited (accession number to follow).

---

**Figure S1.**
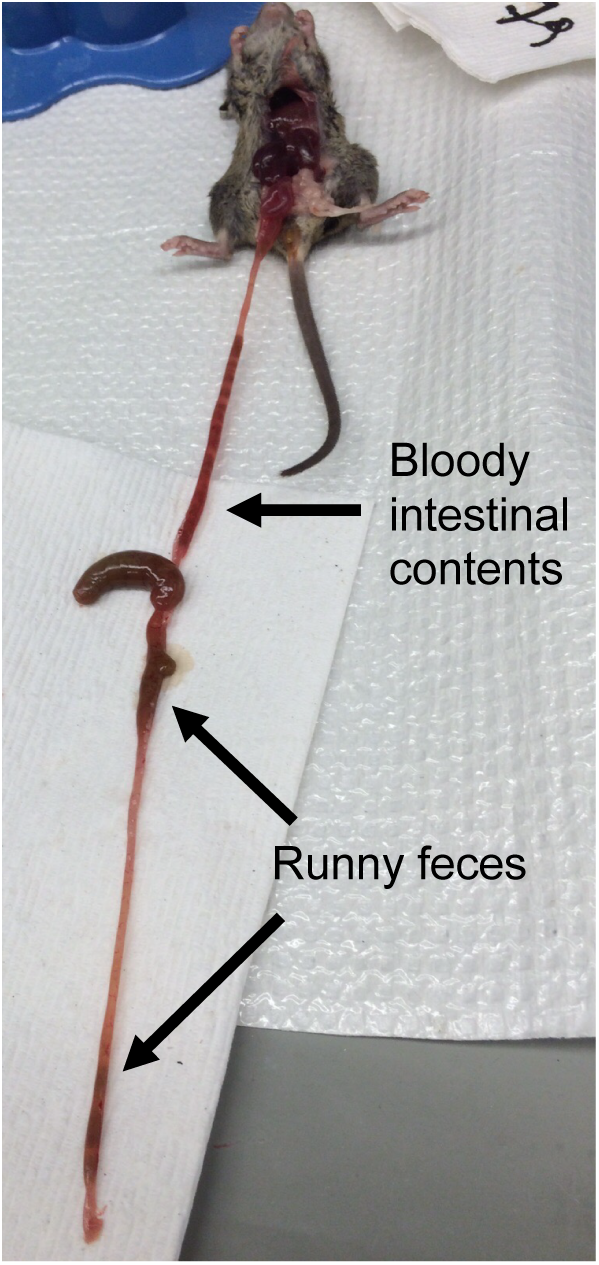
Donor 5-colonized mice exhibit intestinal inflammation. A) A C3H/HeN mouse colonized with the CMA Donor 5 for one week.

**Figure S2.**
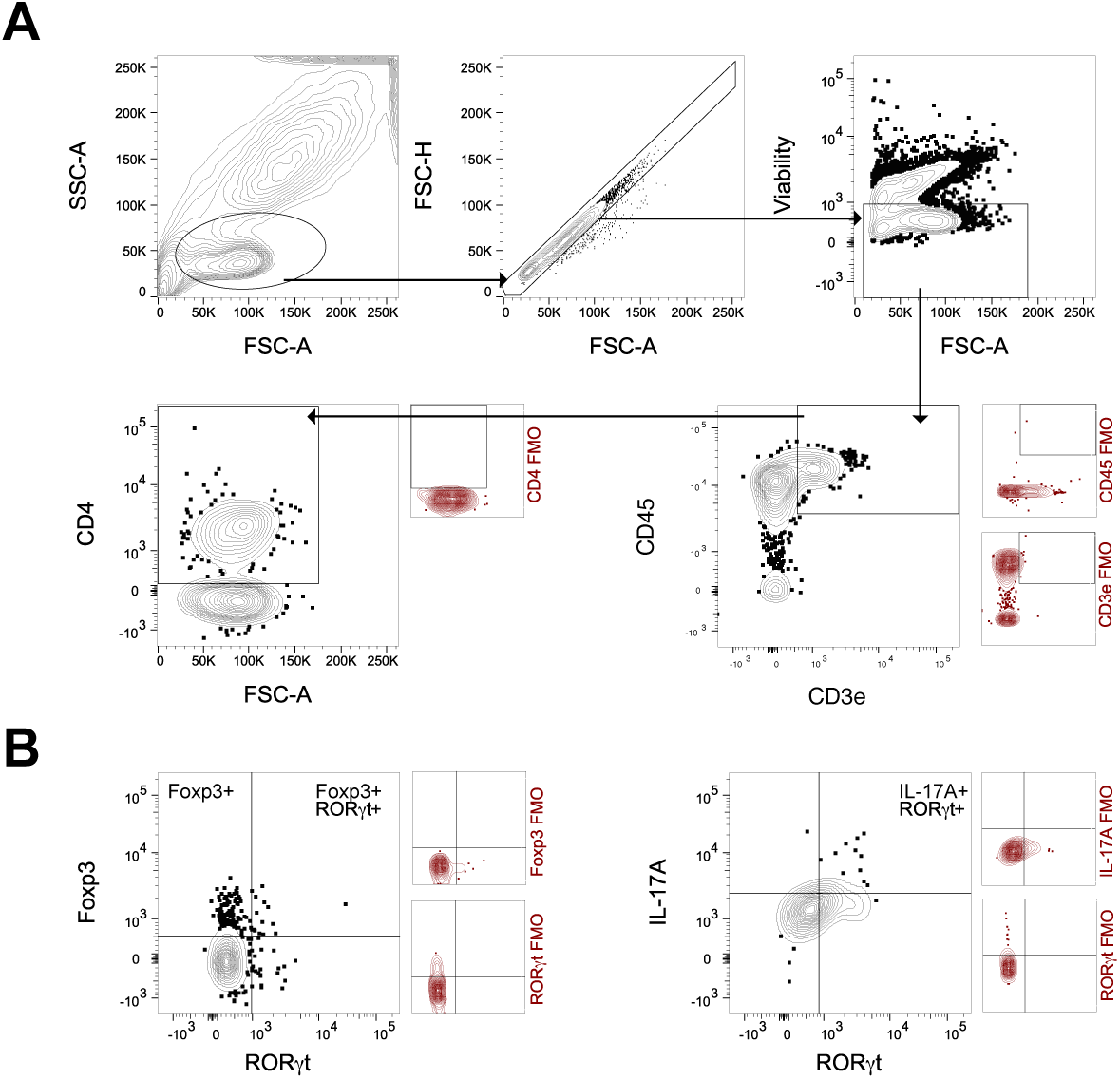
Gating strategy for the identification of Tregs and Th17 cells. A) Gating strategy for the identification of CD4^+^ T cells. B) Gating strategy to identify Foxp3^+^ Tregs, Foxp3^+^ RORγt ^+^ Tregs, and IL-17A^+^ RORγt^+^ Th17 cells within the CD4-positive gate

**Figure S3:**
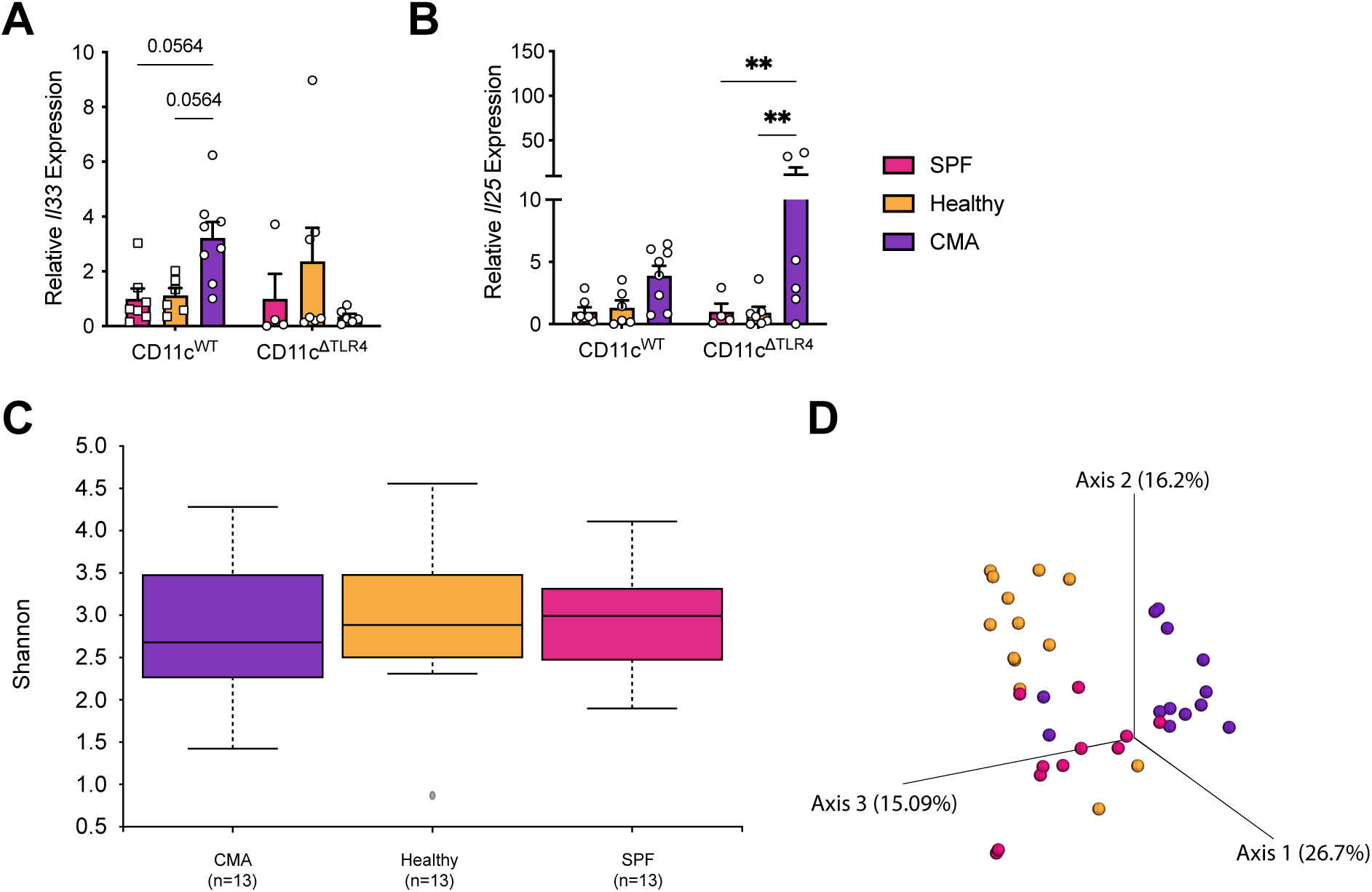
TLR4 signaling in CD11c^+^ cells regulates expression of genes alarmins. Relative expression of A) *Il33* and B) *Il25* in antibiotic-treated SPF mice colonized with the CMA Donor 5 microbiota, healthy Donor 1 microbiota, or left uncolonized after treatment one-week post-weaning/colonization. C) Alpha diversity of each colonization condition (CMA Donor 5, Healthy Donor 1, and SPF, one-week post-colonization/weaning) as measured by the Shannon diversity index D) Principal Coordinate Analysis (PCoA) of beta diversity (Bray Curtis) showing Euclidean distances between antibiotic-treated SPF mice colonized with an infant microbiota (CMA or healthy) or left uncolonized after treatment (SPF).

**Figure S4:**
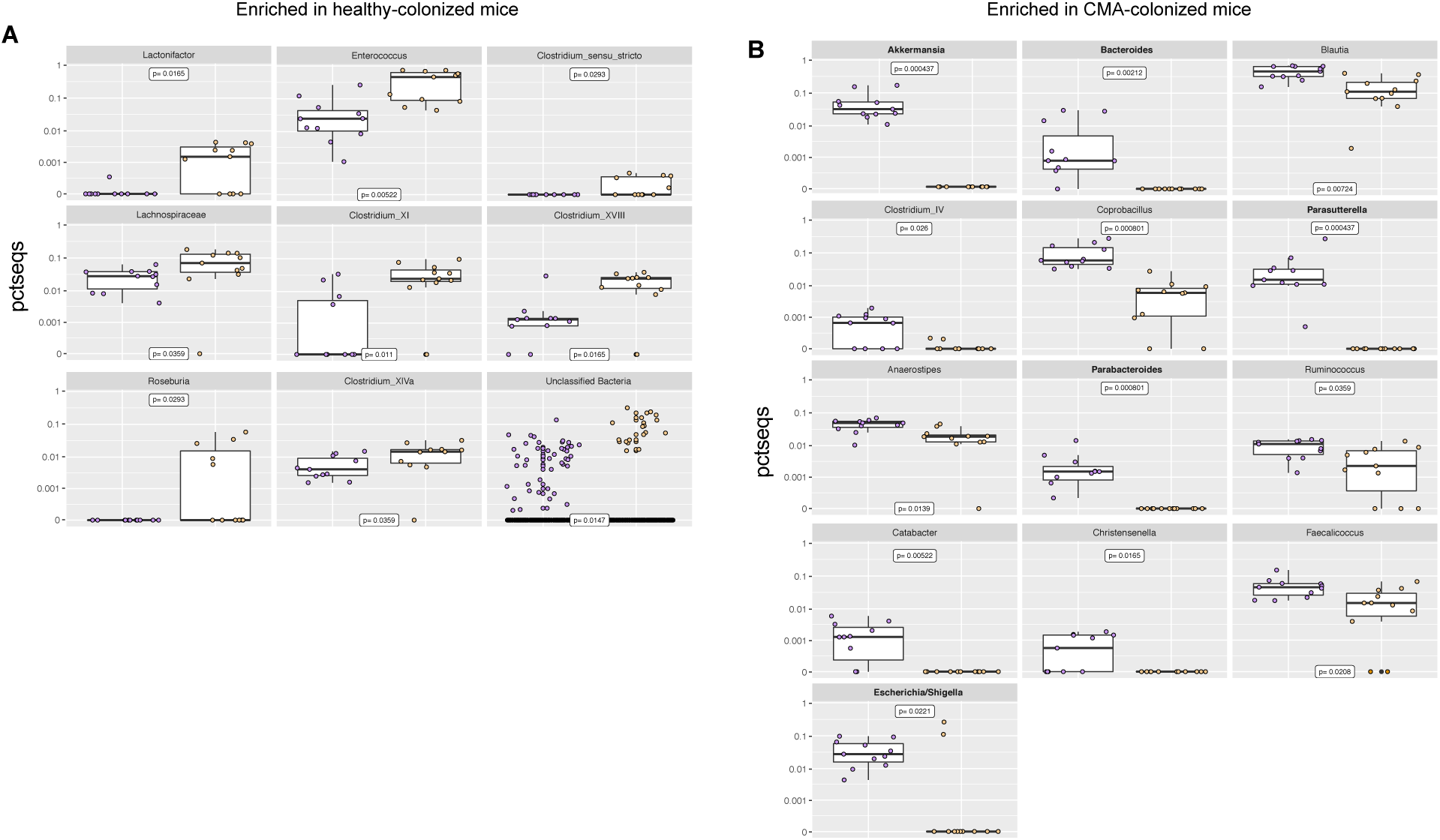
16S Sequencing shows increased abundance of Gram-negative bacteria in CMA- colonized mice. Relative abundance of taxa significantly enriched in A) healthy Donor 2-colonized mice and B) CMA Donor 5-colonized mice one-week post-colonization. Statistical analysis was performed using the Kruskal-Wallis test with a Benjamini-Hochberg correction.

**Figure S5:**
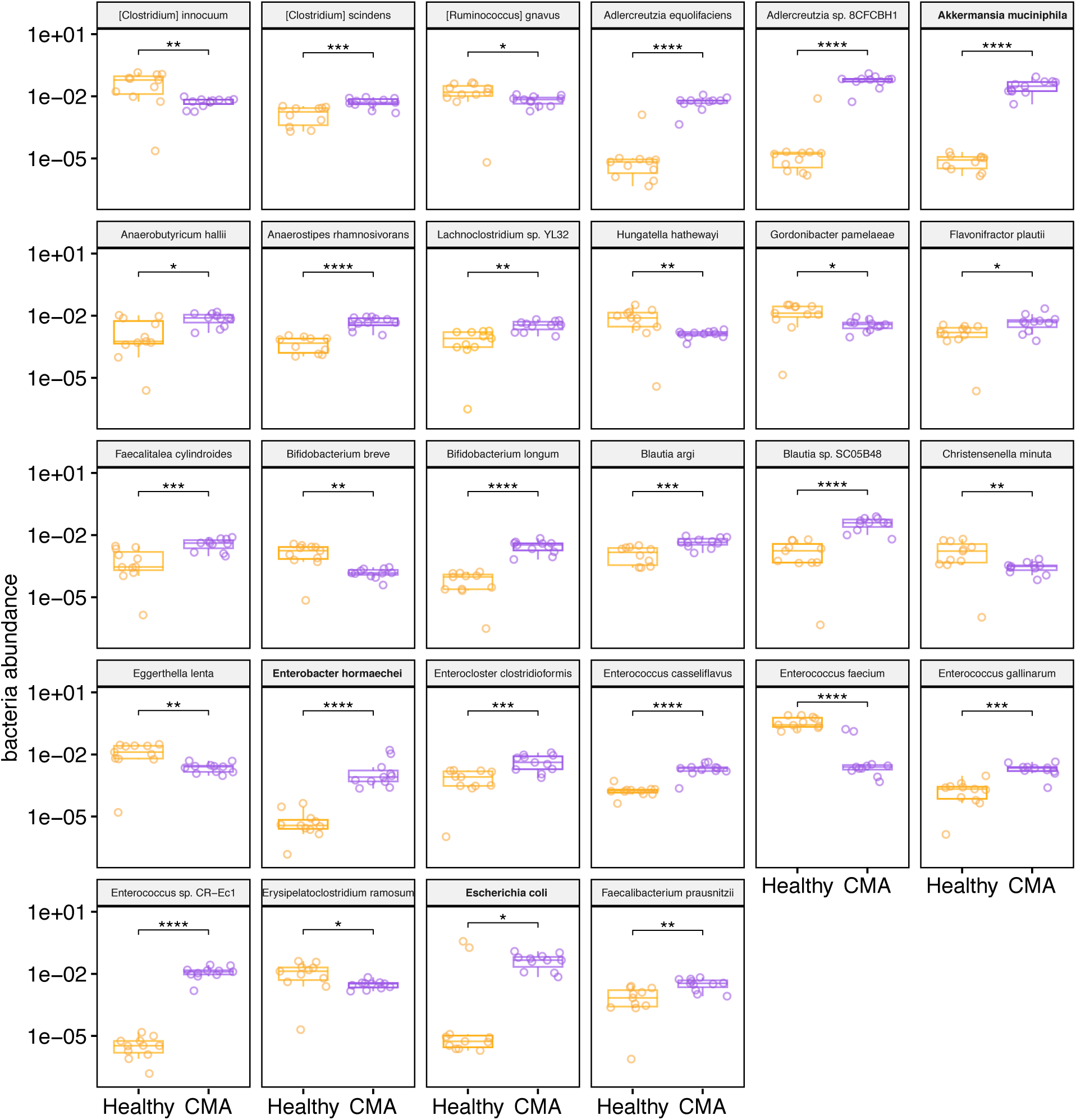
CMA-colonized mice have increased abundance of Gram-negative species, particularly, Bacteroidetes. Differentially abundant bacterial species in healthy Donor 2-colonized and CMA Donor 5- colonized mice one-week post-colonization. Statistical analysis was performed using the Kruskal-Wallis test with a Benjamini-Hochberg correction. \**P* < 0.05, \*\**P* < 0.01, \*\*\**P* < 0.001, \*\*\*\**P* < 0.0001.

**Figure S6:**
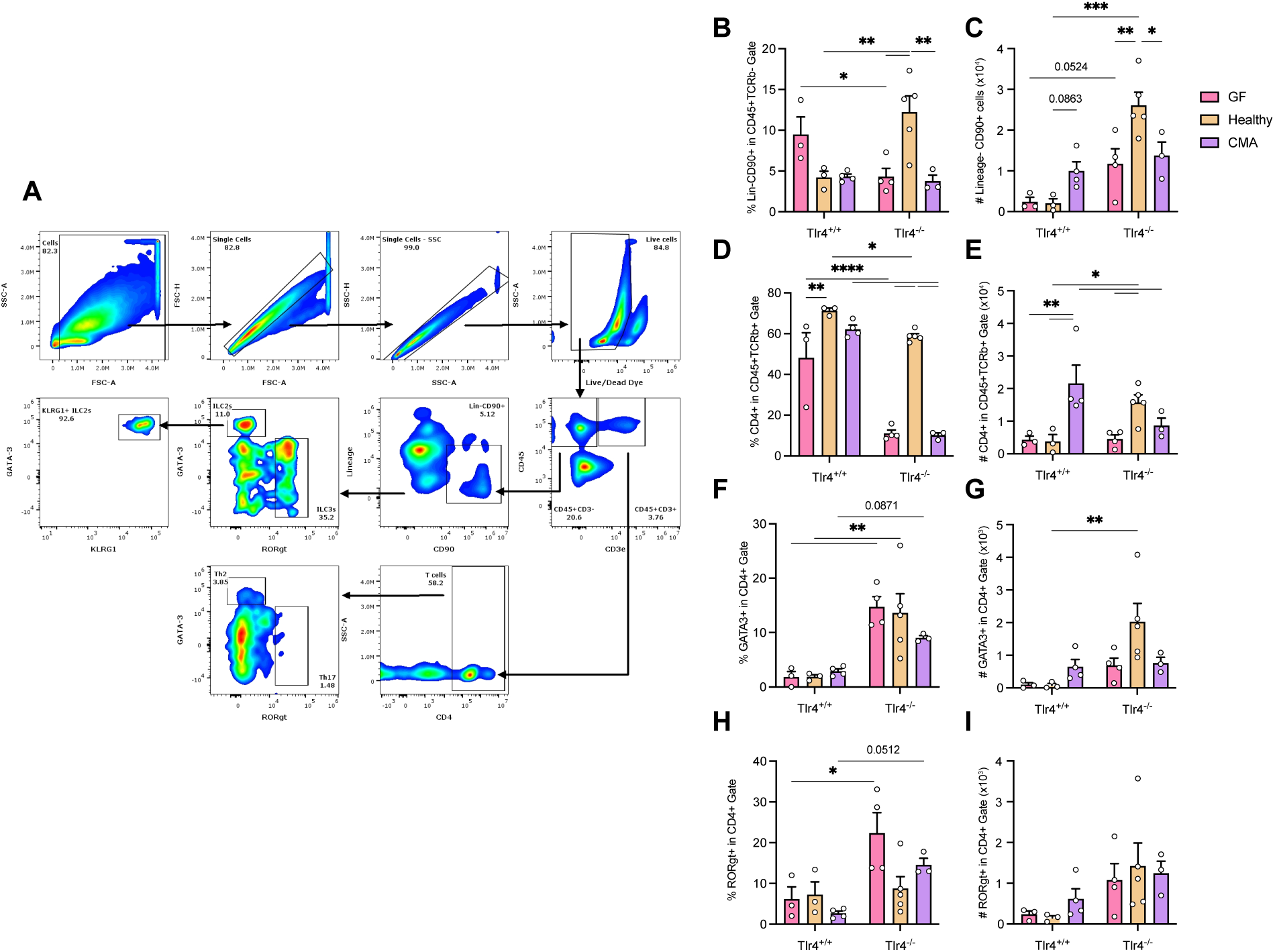
ILC and T cell subsets in sensitized GF, healthy-colonized, and CMA-colonized mice. A) Gating strategy used to identify ILC and T cell subsets in the ileum lamina propria of GF, CMA-colonized, and healthy-colonized mice Frequency and absolute numbers of B-C) total ILCs, D-E) total T cells, F-G) Th2 cells, and H-I) Th17 cells in the ileal tissue of GF, healthy Donor 2-colonized, and CMA Donor 5-colonized *Tlr4*^+/+^ and *Tlr4*^-/-^ mice

## STAR Methods

### LEAD CONTACT AND MATERIALS AVAILABILITY

Further information and requests for resources and reagents should be directed to and will be fulfilled by the lead contact, Cathryn R. Nagler.

#### Recruitment of infant participants

As previously reported (Feehley et al., 2019), fecal samples from healthy infants that had no risk or history of atopic disorders were obtained from participants in a vaccination program at the University of Naples Federico II. Fecal samples from cow’s milk allergic infants were obtained following diagnosis at a tertiary pediatric allergic center (Pediatric Allergy Program at the Department of Translational Medical Science of the University of Naples Federico II); Participation of infants in this study was in compliance with the Declaration of Helsinki and approved by the ethics committee of the University of Naples Federico I. Written informed consent was obtained from the parent(s) or guardian(s) of all children involved.

#### Collection and preparation of human fecal samples

Fresh fecal samples were collected in the clinic in sterile stubs, weighed, and mixed with 2 mL of 30% glycerol in Luria-Bertani broth per 100-500 mg of feces (Feehley et al., 2019). These samples were aliquoted into sterile cryovials and immediately stored at -80 °C. Samples were shipped from Naples, Italy to the University of Chicago on dry ice, where they were stored at -80 °C until homogenization. For colonization of mice, frozen fecal samples were thawed in an anaerobic chamber. Thawed feces were mixed with 3 mm borosilicate glass beads in a sterile 50 mL tube with 2.5 mL of pre-reduced 0.05% Tween in 1X PBS cysteine and vortexed gently to dissociate. The resulting homogenate was filtered through a 100 μm filter for a total of four times. The final filtrate was mixed with an equal volume of 30% glycerol/0.05% cysteine. The solution was aliquoted into Balch tubes with rubber stoppers and introduced into the gnotobiotic isolator. The remaining fecal homogenate was frozen in aliquots at -80 °C.

#### Mouse husbandry and care

All mice used in these experiments were housed and bred at the University of Chicago of Animal Resource Center in a specific-pathogen free (SPF) facility or the Gnotobiotic Research Animal Facility (GRAF) under a 12-hour light/dark cycle. Mouse cages were autoclaved and bedded with pine-shavings. Mice maintained in the GRAF were housed in Trexler-style flexible film isolators (Class Biologically Clean) with Ancare polycarbonate mouse cages (catalog number: N10HT). Gnotobiotic mice were fed an autoclaved diet (5K67, Purina Lab Diets; sterilized at 121 °C for 30 min) and were provided sterile infant formula (Enfamil) or water (USP grade, pH 5.2) *ad libitum* (Feehley et al., 2019). Cage bedding for gnotobiotic mice was changed daily or every other day due to leakage of formula from the bottles.

Sterility of each isolator was confirmed using culture-dependent and culture-independent methods on a weekly basis. Collected fecal pellets were homogenized and cultured in Brain-Heart Infusion, Nutrient, and Sabouraud broth at 37 °C and 42 °C under aerobic and anaerobic conditions, respectively, for 96 hours. DNA isolated from collected fecal pellets was analyzed for the presence of the 16S gene using PCR. Reaction mixtures were made using 40 μL of water, 0.5 μL of dNTP mix, (Thermo Scientific), 1 μL of 8F primer, 1 μL of 1492R primer, 0.5 μL of DreamTaq polymerase (Thermo Scientific), 5 μL of 10X DreamTaq Buffer (Thermo Scientific), and 2 μL of DNA per 50 μL reaction. Cycling conditions for the reaction mixture entailed an activation stage at 95°C for 2 min; 40 cycles of a 30 sec hold at 95°C, a 30 sec hold at 48°C, and a 1 min hold at 72°C; and a final stage at 72°C for 10 min followed by an infinite hold at 4°C. Upon rederivation or acquisition of new animals into the gnotobiotic colony, all mice were screened for parasites, full serology profile, and/or PCR, bacteriology, and gross and histological analysis of major organs through IDEXX Radil or Charles River Lab using an Axenic Profile Screen. Specific-pathogen free mice were fed an irradiated standard chow diet (Teklad 2916, Envigo) and provided autoclaved acidified water *ad libitum*. All experiments were performed in compliance with the University of Chicago Animal Care and Use Protocols (Protocol #72039 and #72114).

#### Mouse lines

Specific-pathogen free C57BL/6 *CD11c*^cre^ and C56BL/6 *Tlr4*^fl/fl^ mice were purchased from Jackson Labs. *CD11c*^cre^ and *Tlr4*^fl/fl^ mice were bred inhouse to obtain a *CD11c*^cre^*Tlr4*^fl/fl^ colony. Specific-pathogen free WT *Foxp3*^GFP^ and *Tlr4*^-/-^*Foxp3*^GFP^mice, both on a C57BL/6J background, were bred in-house. *Foxp3*^GFP^ mice, were crossed with *Tlr4*^-/-^ mice (originally from Jackson Labs) to create a homozygous mutant colony. These mice were rederived by Kathy McCoy at the University of Calgary International Microbiome Center (Calgary, Alberta, CA) by two-cell embryo transfer into germ-free pseudo-pregnant recipient females and bred and maintained in flexible-film isolators. Germ-free C3H/HeN mice were provided by Tatyana Golovkina as an in-house transfer through the GRAF.

#### Colonization of experimental mice with the human infant gut microbiota

Human infant microbiota repository mice were created by gavaging 3-6-week-old C3H/HeN mice with 500 μL of prepared human infant fecal homogenate. The repository mice for each donor were housed in separate isolators to avoid microbial cross-contamination. Fecal samples from both repository and experimental mice were routinely examined through 16S rRNA sequencing analysis, which demonstrated that mouse-to-mouse transfer from repository to experimental mice by gavage was highly reproducible and stable over time. To aid in the transfer of the human infant microbiota, repository mice colonized with the healthy microbiota (Donor 1 or Donor 2) were maintained on Enfamil Infant formula (Mead Johnson Nutrition), while CMA repository mice colonized with the Donor 5 microbiota initially received Nutramigen I formula (Mead Johnson Nutrition) *ad libitum* (that is, the formula the CMA infants consumed) but were switched to Enfamil for the duration of the experiments reported in this thesis.

For colonization experiments, feces from these repository mice were collected and homogenized in 1 mL of sterile PBS. For colonization of germ-free mice with the CMA Donor 5 or healthy Donor 1 or Donor 2 microbiotas, mice were gavaged at weaning with 250 µL of freshly homogenized feces obtained from CMA or healthy repository mice. Prior to colonization, mice were fed Enfamil formula at a 1:4 dilution in water for 4 hours to aid in the transfer of the human infant microbiota. After colonization, CMA-colonized and healthy-colonized mice were housed in specialized isolators, fed standard chow, and were provided with diluted infant formula. On day 7 post-colonization, mice were sacrificed and fecal, cecal, and ileal contents were collected and stored at -20 °C.

For SPF mice, littermates were treated intragastrically with 100 µL of an antibiotic cocktail of kanamycin sulfate (4 mg/mL), gentamycin sulfate (0.35 mg/mL), colistin sulfate (8500 U/mL), metronidazole (2.15 mg/mL), and vancomycin hydrochloride (0.45 mg/mL) daily for 7 days prior to weaning. At weaning, mice were colonized by intragastric gavage with 250 µL of freshly homogenized feces obtained from CMA or healthy repository mice. SPF littermate experimental controls followed the same antibiotic treatment regimen but were left uncolonized at weaning. They were provided sterile acidified water. Mice colonized with the human infant microbiota followed the same experimental timeline as their germ-free counterparts and were sacrificed 7 days post-colonization. Intestinal contents were stored as previously described.

#### Sensitization of mice with β-lactoglobulin (BLG)

At weaning, mice were either colonized with the human infant microbiota or left uncolonized as GF controls. On day 7 post-colonization, mice were fasted for 3 hours. Following this fast, mice were gavaged with 200 µL of sterile 0.2 M sodium bicarbonate in Ultrapure water (Gibco). The mice were allowed to rest for 30 minutes and then each mouse was gavaged with 200 µL of sterile BLG (20 mg) plus cholera toxin (CT, 10µg) in 1X PBS. The food was replaced, and this procedure was repeated on day 14 post-colonization. Mice were sacrificed on day 21 post-colonization.

#### Ileal epithelial cell isolation

At sacrifice, whole terminal ileum tissue (measured 10 cm proximal to the cecum) was carefully removed, being sure to keep the tissue fully intact. Ileal tissues were cleaned, inverted, and inflated with enough air to expand the tissue but not rupture it (Nik et al., 2013). Peyer’s patches were not removed. Intestinal epithelial cells were collected from the inflated inverted tissue in Cell Recovery Solution (Corning) by agitating the tissue every 5 minutes for 30 minutes on ice. Cells were centrifuged at 800 g for 5 min, lysed in 1 mL of TRIzol Reagent (ThermoFisher), and stored at ^-^80 °C for at least 24 hours.

RNA was extracted using the PureLink RNA Mini Kit (Invitrogen). Samples in 1 mL of Trizol were slowly thawed on ice, then incubated at RT for 5 min. Following incubation, 200 lllL of chloroform was added, and each sample was shaken vigorously for 15 seconds. Samples were then incubated at RT for 3 min and centrifuged at 12000 x g for 15 min at 4 °C. Approximately 600 lllL of the colorless upper phase was transferred to a new RNAse-free tube. An equal volume of 70% ethanol was added, and the sample was vortexed. The sample was transferred to a spin column and spun at 12000 x g for 30 sec until the whole sample was processed. The column was washed once with 700 μL of wash buffer (Wash Buffer I), followed by two 500 μL washes of wash buffer containing 70% ethanol (Wash Buffer II). The spin column was centrifuged at 12000 x g for 1 min to dry the membrane. For elution of RNA, 50 μL of RNAse-free water was added to the membrane, and the column was allowed to incubate for 1 minute at RT. The spin column was transferred to a new tube and spun at 12000 x g for 2 min at RT. RNA quality and quantity was immediately determined using Nanodrop One (Thermo Scientific). Samples were stored at ^−^80 °C until utilized for cDNA synthesis or RNA sequencing.

#### Lymphocyte isolation from ileal tissue

Ileal lymphocytes were isolated according to previous protocols, with some modification (Gracz, et al., 2012; Moro et al., 2015). At sacrifice, ileal tissue was cleaned with 1X PBS with mesenteric fat removed and cut longitudinally. The whole tissue was placed into 4 mL of 30 mM EDTA and 1.5 mM dithiothreitol in 1X PBS for 20 minutes on ice. The tissue was then transferred to 4 mL of 30 mM EDTA in 1X PBS and placed into an incubator for 10 minutes at 37 °C, followed by gentle shaking for 30 seconds. Tissues were then transferred to a 6 well plate and washed with 10 ml of 1X PBS to remove remaining epithelial cells. In initial studies for samples being analyzed for T cell subsets, tissues were transferred to a 3 mL digest solution of 4% fetal bovine serum (FBS) in Dulbecco’s Modified Eagle Medium (DMEM) containing 0.5 mg/ml collagenase D (Roche), 0.5 mg/ml dispase (Roche) and 40 μg/ml DNAse (Sigma). In later studies for samples being analyzed for both T cell and innate lymphoid cell (ILC) populations, tissues were transferred to a 3 mL digest solution of 4% FBS in RPMI containing 0.05 mg/mL liberase (Roche) and 40 μg/ml DNAse (Sigma). The tissue was minced finely in the digest solution with surgical scissors and placed into a shaking incubator for 30 minutes at 37°C, 200 rpm.

Following digestion, the cell suspensions were poured over a 70-micrometer cell strainer into a 50 mL conical tube and washed with 20 mL PBS. The cell suspension was spun at 800 g for 5 minutes and resuspended in 4 mL of 4% FBS in DMEM. Four milliliters of 80% Percoll (36 mL of Percoll, 10 mL of 1X PBS, and 4 mL of 10X PBS per 50 mL) was added to the suspension, and the suspension was gently overlaid onto 5 mL of 80% Percoll. The Percoll gradient was centrifuged at 1200 x g for 35 minutes with 0 brake acceleration and 0 brake deceleration. Cells were collected at the interface of the Percoll layers and washed with 30 mL of 4% FBS in DMEM. The cells were then centrifuged at 800g for 5 min, resuspended in cold 4% FBS in DMEM, and counted for viability with trypan blue.

#### Cell stimulation and staining for flow cytometry

For the staining of T cell subsets, ileal lymphocytes from WT and TLR4 KO mice were plated at a concentration of 10^6^ viable cells per milliliter in a stimulation solution containing 50 ng/mL of phorbol myristate acetate (PMA, Sigma) and 750 ng/mL ionomycin (Sigma) in the presence of Brefeldin A (10μg/mL, Sigma). Cells were incubated for four hours at 37°C, 10% CO2. After stimulation, cells were centrifuged at 800 g for 5 minutes and washed in 200 µL of 1X PBS supplemented with 4% FBS for staining in a 96-well round bottom plate. For ILCs, lymphocytes were directly transferred into a 96-well round bottom plate for staining immediately after isolation from ileal tissue and washed with 200 µL of 1X PBS supplemented with 4% FBS.

Following the wash, cells were incubated in Fc block for 10 minutes on ice, followed by surface staining on ice in the dark for 30 min. Cells were washed and then stained for viability 30 minutes on ice in the dark using Live/Dead Fixable Aqua (Life Technologies) at a 1:500 dilution. Cells were then fixed using the Foxp3/Transcription Factor Staining Buffer Set (eBioscience), with prepared solutions according to manufacturer’s protocol, for 40 minutes on ice. Fixed cells were then stained intracellularly for 40 minutes at room temperature in the dark. Data was collected on the Fortessa 4-15 (BD Biosciences) or Aurora spectral flow cytometer (Cytek) and analyzed using FlowJo software (v10.3.2, Becton Dickinson and Company). Antibodies used in these experiments are listed in Table S3.

#### ELISAs

Venous blood samples (3 mL) were collected from CMA and healthy patients at the University of Naples Federico I. Serum was obtained by centrifugation for 15 min and stored at - 80°C. IgE specific for the nBos d 5 epitope on BLG was quantified using the ThermoFisher Scientific ImunnoCAP system and was expressed as kilounits per liter (kUA/L). Serum SAA-1 levels were determined with a Human Serum Amyloid A ELISA Kit (AbCam) with a range of 0.41 ng/mL - 300 ng/mL.

For qualification of fecal IgA, ELISA plates were coated overnight with 2 μg/mL of goat-anti mouse IgA-UNLB (Southern Biotech), 50 μL per well. The next day, plates were washed 4x with 300 μL of 1X PBS-0.05% Tween and blocked with 200 μL of 3% BSA/PBS for at least 2 hours at RT. Fecal pellets were weighed and placed into 0.01% NaN3 in 1X PBS (1 mL/100 mg of feces). Pellets were vortexed for 5 minutes at top speed and centrifuged for 15 minutes at 10,000 rpm. The supernatant was collected and diluted 1:8 or as needed for IgA quantification. Excess sample was stored at -80 °C. A nine-point standard curve was made by conducting serial 1:1 dilutions of 200 ng/mL stock solution of mouse IgA UNLB (BD Pharmigen). Following the 2-hour incubation period, 50 μL of each sample and standard was applied to the wells overnight. The following day, plates were washed 4x with 300 μL of 0.05% Tween in 1X PBS. 50 μL of diluted secondary antibody (1:2500) was applied to each well and incubated for 1 hour at RT. Following incubation, the plate was washed 4x with 300 μL of 0.05% Tween in 1X PBS Tween. Developer (KPL) was applied to the plate, 100 μL per well, and was allowed to develop for up to 30 minutes. Development was stopped with 100 μL per well of 5% EDTA in water.

#### Quantitative Real Time PCR (qRT-PCR)

Complementary DNA was synthesized from isolated RNA using the iScript cDNA Synthesis kit from BioRad using a normalized quantity of RNA for each sample (500ng – 1500ng). Depending on the desired quantity of RNA, the reaction mixture was scaled up or down accordingly. For each 20 μL reaction, 4 μL of 5x Reaction Mix and 1 μL of RT enzyme was used along with an appropriate amount of water depending on sample RNA concentration. Reaction mixtures were placed in a thermocycler and underwent the following conditions to synthesize cDNA: 1) 5 min at 25°C, 2) 20 min at 46°C, 3) 1 min at 95°C, 4) held at 4°C indefinitely. Samples were used right away for qRT-PCR analysis or stored at ^-^20°C for future use.

cDNA reaction mixes were made using the QuantiNova SYBR Green Kit (Qiagen) or PowerUp SYBR Green system (Applied Biosystems). For the QuantiNova Kit, reaction mixtures consisted of 10 μL of 2x SYBR Green PCR Master Mix, 0.1 μL of QN ROX Reference Dye, 1 μL of forward primer, 1 μL of reverse primer, 5.9 μL of RNAse-free water, and 2 μL of cDNA for a total of 20 μL. For the PowerUp kit, reaction mixtures consisted of 10 μL of PowerUP SYBR Green, 1 μL of forward primer, 1 μL of reverse primer, 6 μL of RNAse-free water, and 2 μL of cDNA for a total of 20 μL. Samples were run in duplicate to determine the average cycle for which each sample reached a set threshold (0.2). Primers used in the reaction mixes are listed in Table S3. Expression was measured with the QuantStudio 3 instrument by Applied Biosystems. The cycling conditions for the reaction mixture entailed an activation cycle of 50°C for 2 min followed by one cycle of 95°C for 10 min and 40 cycles at 94°C for 20 s, 55 °C for 20 s and 72 °C for 50 s. The fluorescent probe was detected in the last step of the cycle. A melt curve was performed at the end of the PCR reaction to confirm the specificity of the PCR product. Relative expression was calculated by determining the ΔCt based on expression of the endogenous housekeeping gene (*Hprt*), followed by normalization to the control group (GF or antibiotic-treated SPF) within each genotype.

#### Bulk RNA sequencing and analysis

RNA libraries for total RNA from ileal epithelial cells were prepared and sequenced at the University of Chicago Functional Genomics Core. Libraries were prepared using a TruSeq Stranded Total Library Preparation Kit with Ribo-Zero Human/Mouse/Rat (Illumina). Samples were sequenced using 50-bp single-end reads chemistry on the HiSeq2500 instrument. Raw reads were assessed for quality, counted, and aligned to the mm10 reference genome using the R- subread package (Liao et al., 2019). All samples had a Phred score of at least 30. Normalization and differential expression analysis between healthy- and CMA-colonized were performed using the edgeR package (Chen et al., 2016). Read counts for each gene were fit to a negative binomial generalized log-linear model. Multiplicity correction was performed by applying the Benjamini-Hochberg method on the *P*-values to control for the false discovery rate (FDR = 0.1). Genes with an FDR of <0.05 and a fold change >1.5 or <-1.5 were plotted as significant. Gene ontology pathways for genes expressed in CD11c^WT^ and CD11c^ΔTLR4^ CMA-colonized mice were determined using Metascape (Zhou et al., 2019).

#### DNA extraction from fecal and ileal contents

DNA was extracted using the QIAamp PowerFecal Pro DNA kit (Qiagen). Samples were suspended in PowerBead tubes along with lysis buffer and loaded on a Bead Mill 24 Homogenizer (Thermo Fisher) or taped to a vortex. Tubes were agitated at maximum speed for 10 minutes at RT to dissociate the fecal pellet. PowerBead tubes were then centrifuged at 10,000 x g for 30 seconds at RT. Isolation of released DNA was then conducted according to the manufacturer’s protocol. DNA was purified routinely using a spin column filter membrane and quantified using the Qubit 4 Fluorometer (Thermo Fisher). Ileal contents were subjected to DNA isolation using the PowerSoil® DNA Isolation Kit (MoBio) or the QIAamp PowerFecal Pro DNA kit (Qiagen), according to the manufacturer’s protocol.

#### 16S Sequencing and Analysis

Isolated DNA was sequenced at the Argonne National Laboratory Sequencing or the Duchossois Family Institute (DFI) at the University of Chicago. For samples sequenced at Argonne, 16S rRNA gene amplicon sequencing was performed on an Illumina MiSeq instrument. Paired-end reads (151 bp) were generated with 12-bp barcodes. The V4 region of the 16S rRNA gene was amplified using 515F and 806R primers that included sequencer adapter sequences used in the flow cell. Raw multiplexed reads were provided by the facility and were demultiplexed using qiime2 (v.2018.4, Bolyen E et al., 2019).

For samples sequenced at the Duchossois Family Institute, sequencing was performed on the Illumina MiSeq platform in the Functional Genomics Facility at the University of Chicago using 2×250 Paired End reads, generating 5,000-10,000 reads per sample. The V4-V5 region within 16S rRNA gene was amplified using the universal bacterial primers – 563F (5’-nnnnnnnn-NNNNNNNNNNNN-AYTGGGYDTAAA- GNG-3’) and 926R (5’-nnnnnnnn-NNNNNNNNNNNN-CCGTCAATTYHT- TTRAGT-3’), where ‘N’ represents the barcodes, ‘n’ are additional nucleotides added to offset primer sequencing. Amplicons were then purified using a spin column-based method (Qiagen), quantified, and pooled at equimolar concentrations. Illumina sequencing-compatible Unique Dual Index (UDI) adapters were ligated onto the pools using the QIAseq 1- step amplicon library kit (Qiagen). Library quality control was performed using Qubit 4 Fluorometer (Thermo Fisher) and Tapestation (Agilent) and sequenced on Illumina MiSeq platform to generate 2x250bp reads.

For 16S analysis, Dada2 (v1.14, Callahan, et al., 2016) was used for processing raw 16S rRNA reads. The first 180 bp for both forward and reverse reads were trimmed to remove low quality nucleotides. Taxonomic analysis was determined using the RDP database (Wang et al., 2007) with a minimum bootstrap confidence score of 50. Multiple sequence alignment was performed using msa (v1.18.0, Bodenhofer et al., 2015) and ape (v5.3). Alpha and beta diversity were calculated using Dada2.

#### Shotgun metagenomics

Libraries were prepared using 100 ng of genomic DNA using the QIAseq FX DNA library kit (Qiagen). DNA was fragmented enzymatically into smaller fragments and desired insert size was achieved by adjusting fragmentation conditions. Fragmented DNA was end repaired and polyA tails were added to the 3’ ends to stage inserts for ligation. During the ligation step, Illumina compatible Unique Dual Index (UDI) adapters were added to the inserts and the prepared library was PCR amplified. Amplified libraries were cleaned up, and QC was performed using a Tapestation (Agilent). Sequencing runs were performed on the Illumina NextSeq platform in the Functional Genomics Facility at at the University of Chicago using the 2×150 paired-end reads cassette.

For metagenomic analysis, quality reads were used in Kraken2 (v.2.1.1) for taxonomic classification and to assemble contigs via Megahit (v1.2.9, Li et al., 2015). Genes were annotated using Prodigal (v.2.6.3, Hyatt et al., 2010), and functional annotations were perform using eggnog-emapper (v.2.0.1b, Cantalapiedra et al., 2021). Wilcoxon rank sum test was performed on MAGs identified in healthy-colonized and CMA-colonized mice to determine differentially abundant species.

#### Statistical Analysis

Statistical analyses were performed using GraphPad Prism version 9.3.0. The test details are indicated in the figure legends.

---

**Table 1:**
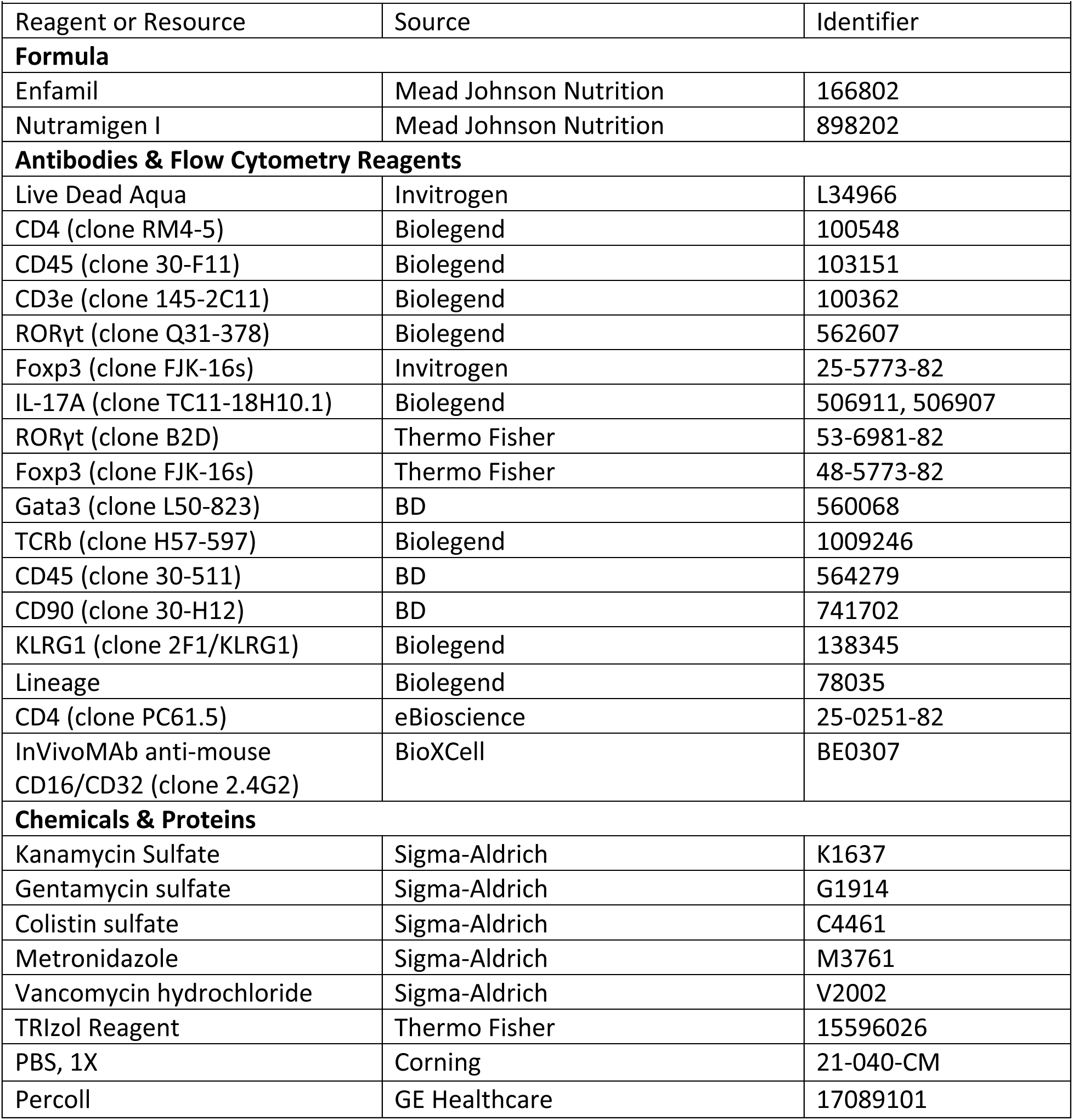
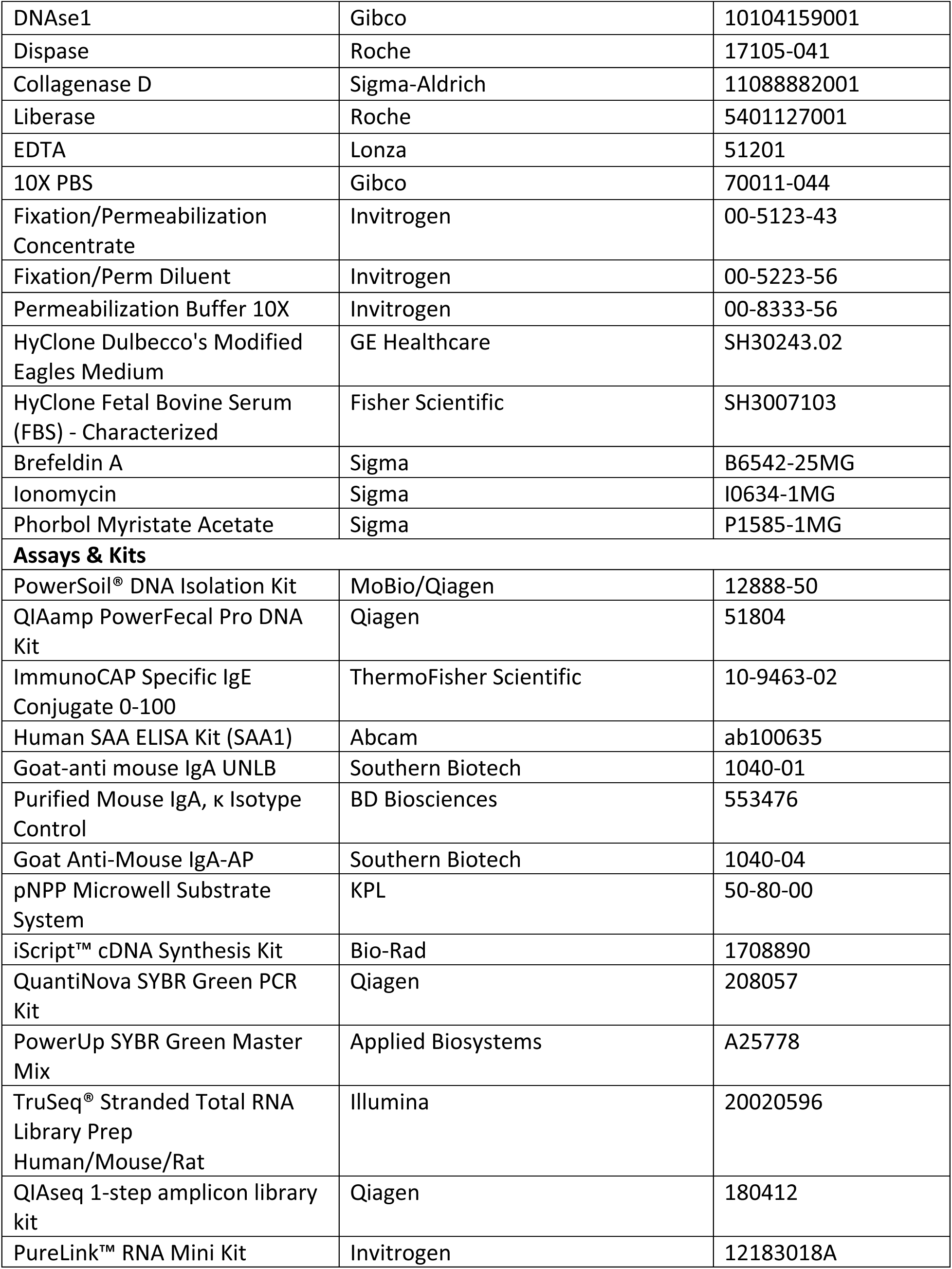
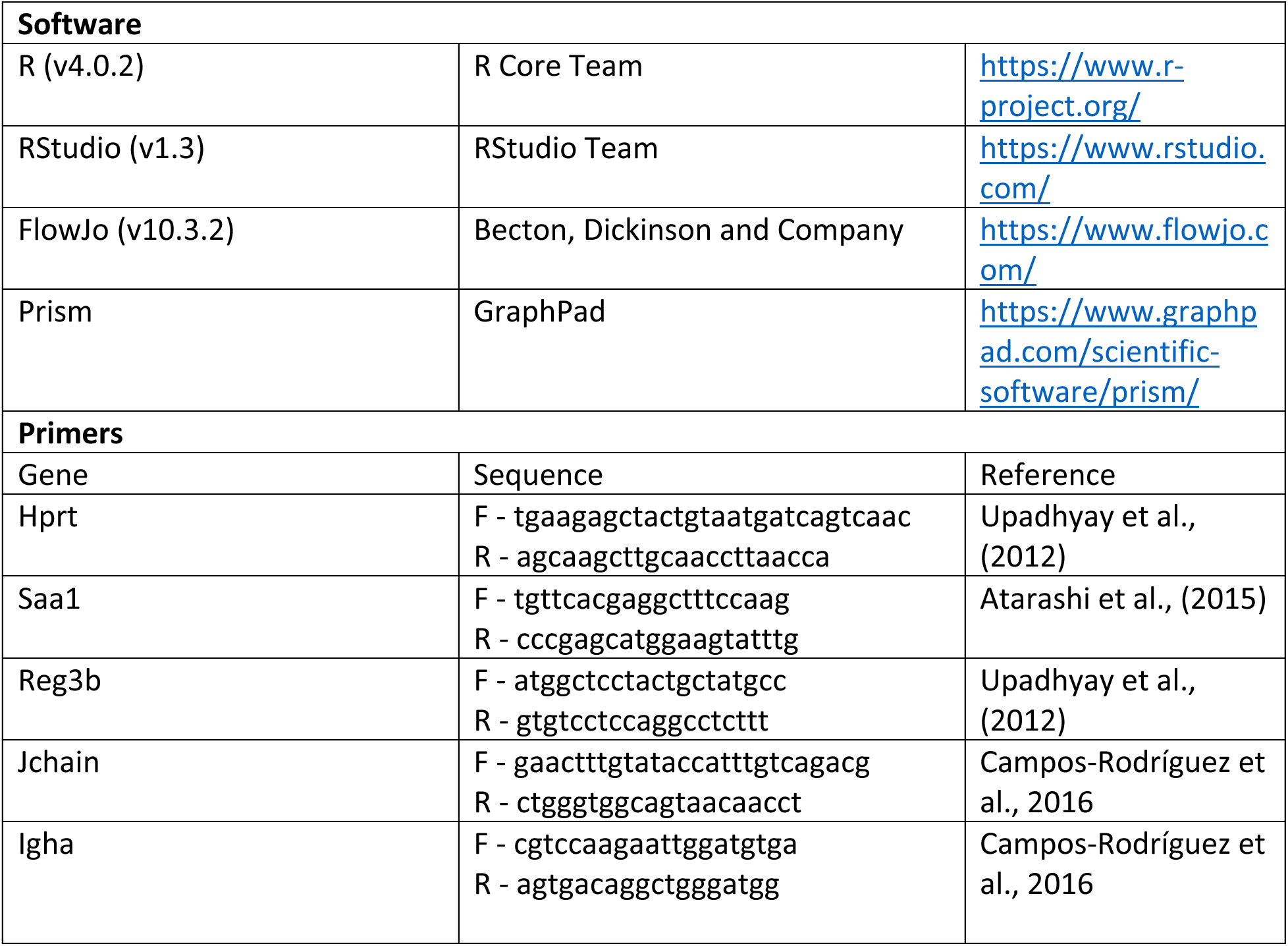
Reagents and resources used in this study

